# Single nucleus RNA sequencing and spatial transcriptomics reveal unique functionalities of gray matter versus white matter oligodendrocytes in aging and Alzheimer’s Disease

**DOI:** 10.64898/2026.08.29.747923

**Authors:** Joseph P. Voth, Javier A. Ramos Benitez, Angela M. Wilson, Scott R. Kennedy, Mamatha Damodarasamy, Amanda Kirkland, Jeremy A. Miller, Caitlin S. Latimer, Emily Ragaglia, Amber Nolan, Daniel D. Child, Lisa M. Keene, Cindy L. Reichel, Brendan F. Kohrn, Larry S. Sherman, Stephen A. Back, C. Dirk Keene

**Affiliations:** Graduate Program in Neuroscience, University of Washington School of Medicine; Medical Scientist Training Program, University of Washington School of Medicine; Department of Laboratory Medicine and Pathology, University of Washington School of Medicine; Allen Institute, Seattle, Washington; Division of Neuroscience, Oregon National Primate Research Center, Oregon Health & Science University; Department of Pediatrics and Neurology, Oregon Health & Sciences University; Department of Pathology, University of California San Diego

## Abstract

Oligodendrocyte (OL) dysfunction and white-matter (WM) vulnerability are increasingly recognized as important aspects of aging and Alzheimer’s Disease (AD), yet human WM–focused, cellular-resolution transcriptomic data remain limited. Here, we profiled prefrontal WM from 48 brain donors spanning young adulthood and late-life with low versus high AD Neuropathologic Change (ADNC) using single-nucleus RNA sequencing followed by spatial transcriptomics (CosMx) in a subset of matched donors. We integrated aged WM OLs with a reference dorsolateral prefrontal cortex grey-matter (GM) OL dataset (SEA-AD) to define region- and pathology-associated OL programs. Across modalities, GM OLs exhibited a robust synapse/neurotransmitter-associated transcriptional signature relative to WM OLs, whereas this program was reduced with aging and attenuated in high ADNC GM. In contrast, WM OLs showed stronger immune-associated programs with aging and further enhancement in high ADNC, including cytokine/chemokine signaling and antigen presentation–related pathways. High ADNC WM OLs also displayed amplified proteostasis and stress-adaptation signatures, including selective upregulation of chaperone/heat shock genes and ferritin subunits, consistent with increased protein-folding demand and altered iron handling. To resolve OL state organization beyond static differential expression, we annotated OL sub-states using marker panels and inferred pseudotime-guided directional state-to-state flows within each tissue/condition stratum. This analysis identified a conserved newly formed differentiating (NFOL)/differentiating → lipid remodeling (APOE/ABCA1/LPL+) → Stress/ISR-reactive architecture, with a pronounced expansion of the Stress/ISR-reactive compartment and altered transition-associated pathway enrichment in high ADNC WM. Together, these data define WM-specific OL programs linked to aging and ADNC and nominate a stress/immune-enriched OL state landscape consistent with a putative senescence-like phenotype in diseased WM.

## Introduction

Oligodendrocytes (OLs) are the myelinating glial cells of the central nervous system and are essential for saltatory conduction along axons [73, 81]. OLs also intimately sustain the neurons they ensheathe by providing trophic support [27, 28], shuttling lactate to axons (via monocarboxylate transporter 1 [30, 50]), and directly stimulating axon mitochondria to increase energy production via extracellular vesicular delivery of SIRT2 [24]. As such, OLs are active participants and crucial for proper nervous system function. Recent studies have identified white matter (WM) reduction as an early signature of Alzheimer’s Disease (AD) [23, 48] underscoring the need to understand how WM oligodendrocytes contribute to AD pathophysiology and progression. While numerous single cell RNA sequencing studies have profiled cortical and mesial temporal brain regions in AD [31, 36, 55, 56], comparatively fewer datasets focus specifically on human WM. Defining disease-associated cell phenotypes and pathways within the transcriptional landscape of human WM OLs in the context of AD can help identify novel therapeutic targets and inform future diagnostic strategies.

Although OLs are distributed throughout the brain, including in grey matter (GM), our understanding of the differences between GM and WM OLs is limited. WM OLs typically generate concentric, tightly compacted myelin sheaths of consistent thickness with equally spaced nodes of Ranvier, whereas GM OLs often form single, partial, or loosely wrapped myelin segments, sometimes with substantial gaps between wraps [21, 86]. Given the significant differences in cytoarchitecture, neuronal activity, and metabolic demand between GM and WM, a clearer understanding of OL state differences between these regions is necessary to properly interpret age- and disease-related changes.

Cellular senescence, classically defined as a state of permanent cell cycle arrest in response to cellular stress or damage [37], has been implicated in multiple cell types in AD mouse models including astrocytes [16], microglia [69], and potentially oligodendrocyte precursor cells (OPCs) [34, 90]. Senolytic therapies have reversed cognitive decline in genetic mouse models of AD [19, 90] and at least one clinical trial evaluating dasatinib and quercetin is currently underway [35, 51]. However, the specific brain cell types targeted by senolytic approaches remain unclear. Although senescence is classically associated with proliferative cell types, recent studies have expanded this definition to include any terminally differentiated cells that adopt aberrant, stress-associated functional states [39, 67]. OLs may be particularly vulnerable to such states given that they produce massive amounts of cholesterol and lipids *de novo* [11], maintain the highest iron content among neural cell types [71] and experience substantial oxidative and metabolic stress [11]. Senescence-like programs, such as activation of DNA damage responses, unfolded protein response pathways, metabolic stress, aberrant immune functions signaling, and cell cycle inhibition, have also been described in glia progenitors [34]. These observations raise the possibility that human OLs adopt durable stress-associated or senescence-like phenotypes during aging and AD.

Here, we sought to determine whether a senescence-like state is present in human WM OLs in the context of aging and AD neuropathology. To address this question, we developed a WM-focused single cell and spatial transcriptomic profiling strategy to phenotype OLs from young adults and elderly individuals, with or without AD neuropathologic change (ADNC). For reference and to understand WM-specific differences, we compared these WM OLs to GM OLs derived from the Seattle Alzheimer’s Disease Brain Cell Atlas (SEA-AD) dataset [31], which includes individuals over 70 years of age from the same biorepository and using identical inclusion and exclusion criteria [31]. Eighteen donors were included in both cohorts, providing a means to profile OL across and within donors. Through these analyses, we define distinct characteristics of WM and GM OL populations and identify a putative senescence-like phenotype enriched for WM OLs in the setting of ADNC.

## Results

### snRNA sequencing and spatial transcriptomics in WM OLs

To better understand the pathophysiology of WM changes in aging and in association with AD, we used single nucleus and spatial transcriptomic profiling specifically focused on characterizing cell type variation and pathway differences. We selected a cohort of brain donors over the age of 70 with high ADNC [44] (n=19); for controls, we developed a cohort of donors with “not” or “low” ADNC that were matched by age, sex, year of death, post-mortem interval, and comorbid neuropathologies (n=13). The ADNC metric is a standard diagnostic format for AD pathological burden across a four-level composite scale of AD (not, low intermediate, and high ADNC) that takes into account beta amyloid plaque distribution (Thal phase), neurofibrillary tangle distribution (Braak stage), and cortical neuritic plaque density (CERAD score) [44]. To understand the impact of age on WM OL lineage, we included a cohort of younger donors between the ages of 27-50 years (n=16). This age-range was selected to minimize confounding effects associated with developmental myelination, which largely concludes in early adulthood [78, 80], and to remove the potential for impact of preclinical or incipient neurodegenerative pathologies (**Population summary statistics in Supplemental Table 1**). To limit regional variability, all samples were obtained from the prefrontal white matter approximately 1 cm anterior to the frontal pole of the lateral ventricle, deep to the cortex and excluding subcortical U fibers.

**Supplemental Table 1.** Cohort demographics and neuropathologic summary statistics. Summary of cohort composition across WM snRNA-seq donors, SEA-AD DLPFC GM reference donors, and CosMx donors, including sex distribution, age at death, and Alzheimer’s neuropathologic change (ADNC) components (Thal phase, Braak stage, CERAD), as well as comorbidity measures (e.g., CAA score, microinfarcts), reported as group-level summary statistics (with ranges where applicable).

| Group | Sex | Age at Death | ADNC Rank | Thal Phase | Braak Stage | CAA Score | CERAD | Microinfarcts |
| --- | --- | --- | --- | --- | --- | --- | --- | --- |
| Young WM | 6F/10M | 39 (27-50) | 0.067 (0-1) | 0.125 (0-1) | 0 | 0 | 0 | 0 |
| Low AD WM | 8F/5M | 87.7 (72-102) | 0.54 (0-1) | 0.77 (0-2) | 2.77 (0-4) | 0.46 (0-2) | 0.23 (0-1) | 0.69 (0-2) |
| High AD WM | 11F/8M | 86.6 (73-99) | 3 (3) | 4.63 (4-5) | 5.47 (5-6) | 1.26 (0-2) | 2.68 (2-3) | 3.74 (0-26) |
| Allen Institute DLPFC GM Low AD | 24F/17M | 89.76 (72-102) | 1.61 (0-3) | 2.5 (0-5) | 4 (0-6) | 0.92 (0-2) | 1.24 (0-3) | 1.02 (0-8) |
| Allen Institute DLPFC GM High AD | 24F/15M | 87.69 (70-99) | 2.64 (1-3) | 4.17 (1-5) | 5 (2-6) | 1.24 (0-3) | 2.43 (0-3) | 0.95 (0-10) |
| CosMx – Low AD | 2F/1M | 86, 87, 98 | 0.33 (0-1) | 4.33 (4-5) | 5.33 (5-6) | 0.66( 0-2) | 0.33 (0-1) | 0.33 (0-1) |
| CosMx – High AD | 2F/1M | 84, 83, 98 | 3 | 1 (0-2) | 3 (2-4) | 1 (0-2) | 2.67 (2-3) | 5.67 (1-8) |

**Supplemental figure 1a-c** displays the process from WM samples to the final sequencing data. Following integration of all WM samples, we used candidate genes to determine general cell type identity for each cluster. While identifying cell types, there was typically one cluster that represented “true” cells and a second cluster that localized between this “true” cluster and the main OL cell grouping and expressed elevated MBP/MOBP. These “smear” clusters that expressed both myelin transcripts and other transcripts unique to other cell types were suspected to be doublets. A doublet is a technical limitation of the emulsion-based single cell transcriptomic procedure where an individual barcode labels two separate nuclei as one nucleus. For our 8,000 nuclei capture target, 10x Genomics estimates a 6.4% doublet rate, while our “smear” clusters represented 8.8% of cells captured (**Supplemental Figure 2**). Individual cluster identities were discovered using two candidate genes’ expression representing each broad cell type (**Fig. 1a**). The resulting UMAP following doublet removal is shown in **Fig. 1b**. In each donor, myelinating OLs were the most frequent cell type captured with 77.3% OLs in high ADNC, 76.75% in low ADNC, and 76.64% in young WM samples (**Fig. 1c**).

**Supplemental Figure 1.**
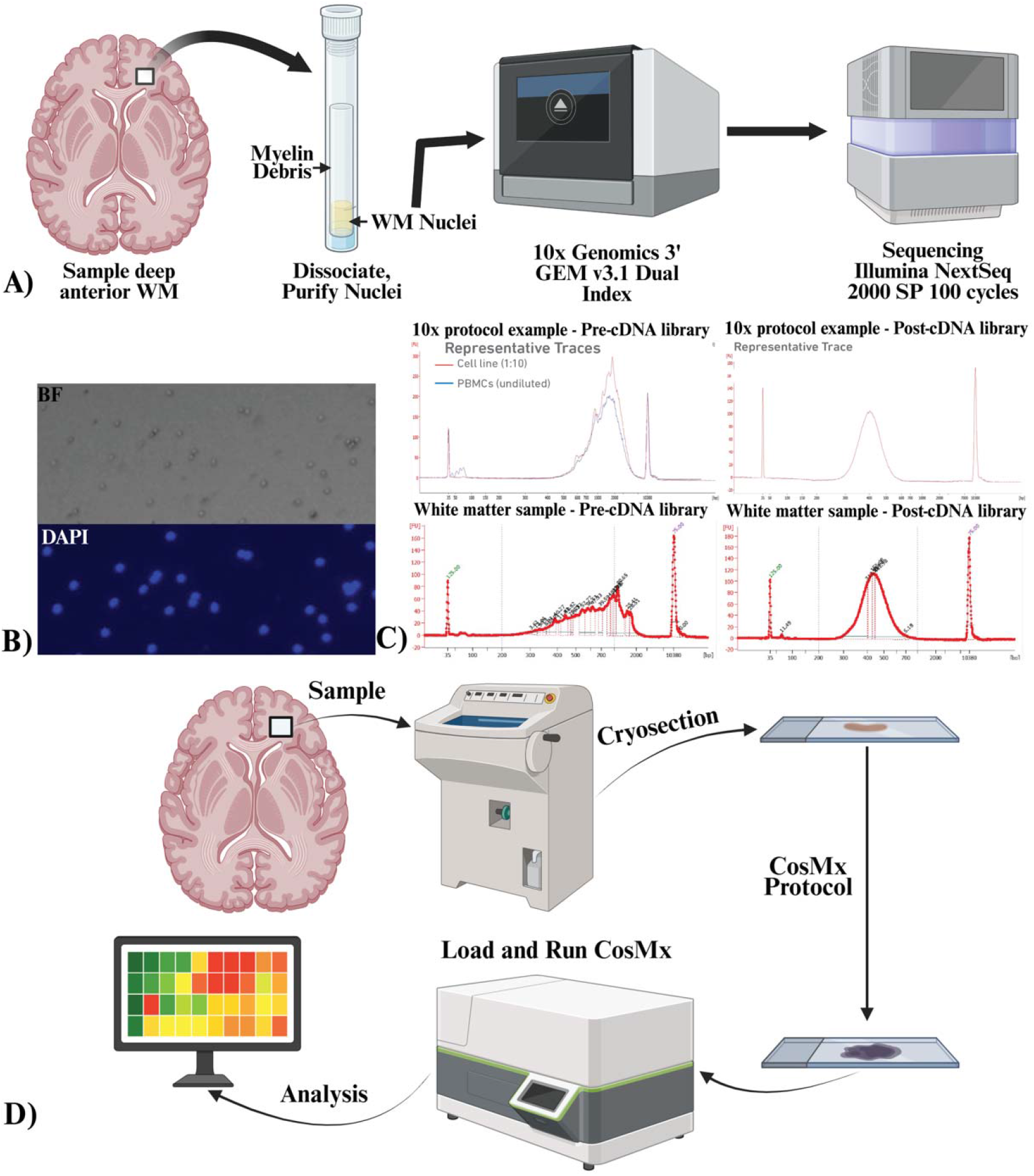
Experimental workflow for WM snRNA-seq and spatial transcriptomics. **(A)** Overview of the white matter (WM) single-nucleus RNA-seq workflow: deep anterior WM sampling, nuclei dissociation and purification (including removal of myelin debris), 10x Genomics Single Cell 3′ (v3.1 dual index) library preparation, and Illumina sequencing. **(B)** Representative images of purified nuclei following isolation, shown in brightfield (BF) and DAPI to illustrate nuclei integrity and sample purity prior to 10x loading. **(C)** Representative Bioanalyzer traces illustrating library QC: pre-cDNA and post-cDNA traces from a representative WM run, used to assess fragment size distributions and guide PCR cycle selection based on quantified cDNA peak profiles. **(D)** Overview of the CosMx spatial transcriptomics workflow: sampling from the targeted region, cryosectioning, CosMx assay execution, and downstream computational analysis.

**Supplemental Figure 2.**
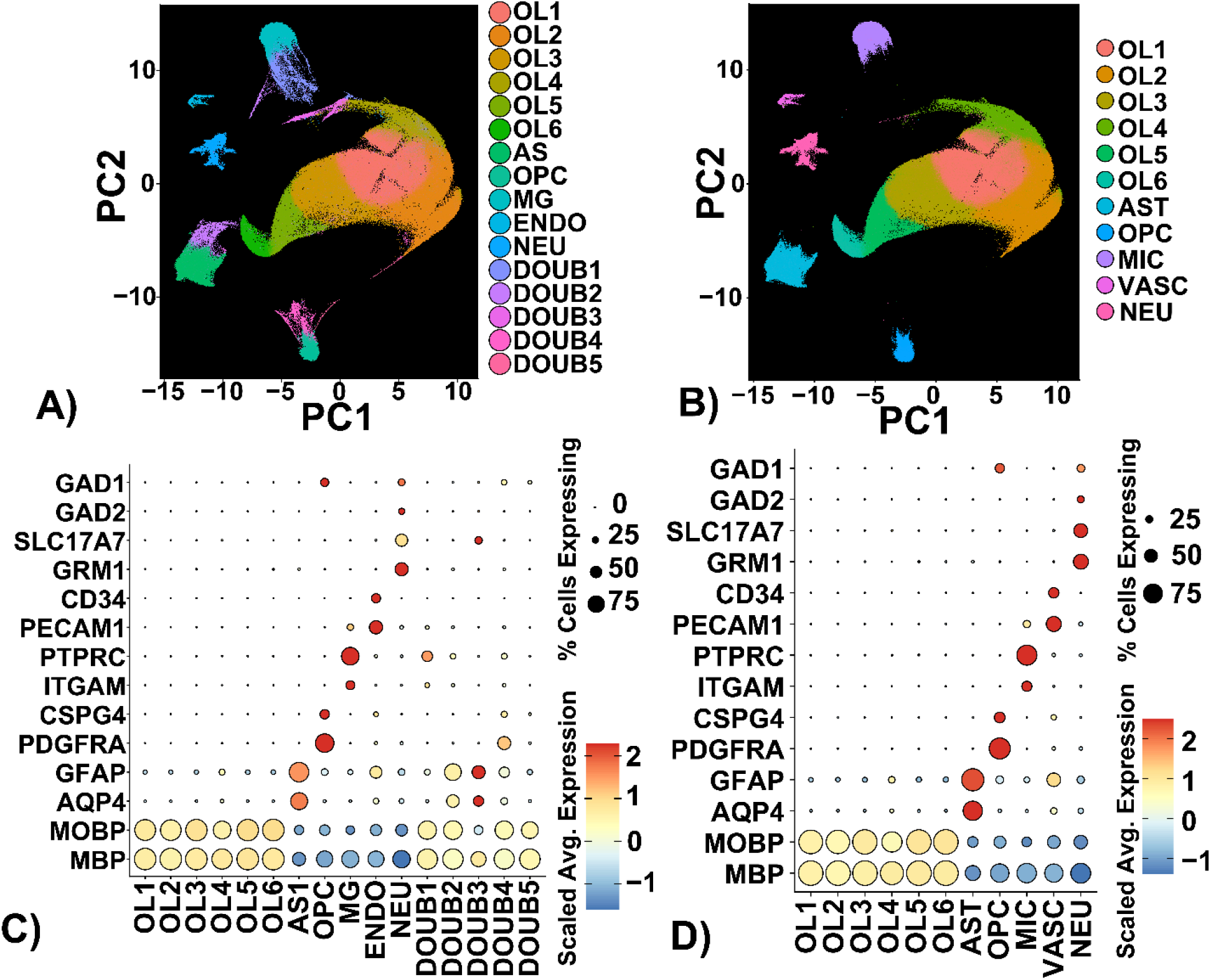
Identification of doublet (“smear”) clusters during WM cell type annotation. A) UMAP of integrated WM nuclei colored by broad cell class assignments based on canonical marker genes. In addition to “true” cell-type clusters, a subset of nuclei form intermediate “smear” clusters positioned between major cell-type groupings and the dominant oligodendroglial cluster, characterized by elevated myelin transcripts (e.g., MBP/MOBP) together with non-oligodendroglial features (C). These intermediate clusters were interpreted as likely doublets (technical multiplets captured under a single barcode), consistent with expected multiplet rates for 10x Genomics droplet-based capture and were flagged and removed for downstream analyses (B and D).

**Figure 1.**
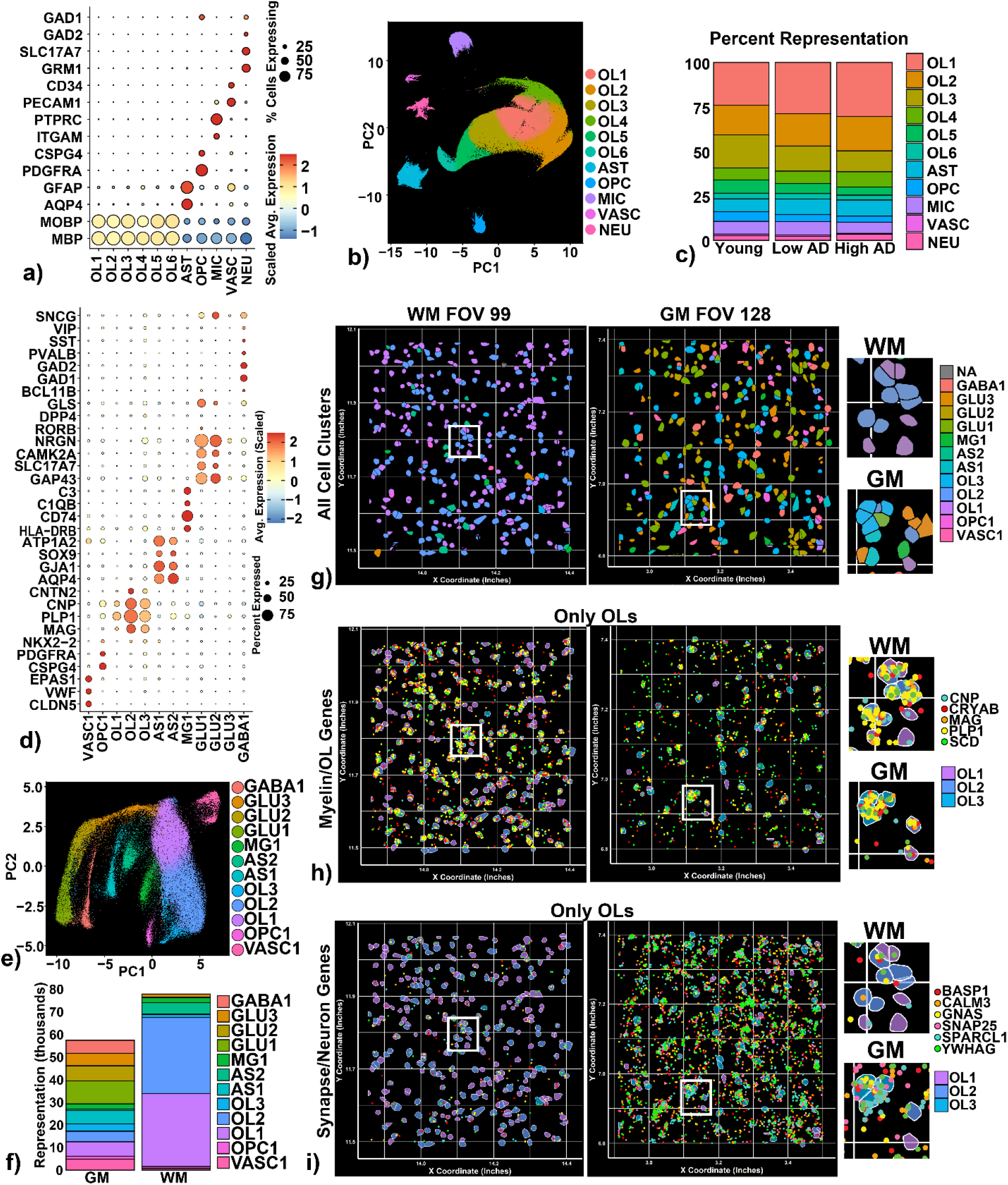
Characterization of the single cell RNA sequencing and CosMx datasets of human brain. **a)** Dot plot of each cluster and its expression across a list of candidate genes used to identify cell types within the snRNAseq. WM dataset. Circle size indicates the percentage of cells within the cluster expressing each gene of interest while dot color represents its scaled expression relative to all other clusters. A stronger red hue indicates this cluster expressed relatively more of that transcript than all others, while blue hues represent relatively less expression. **b)** UMAP following CCA-integration of all individual snRNAseq. WM samples collected show all clusters identified following k-nearest neighbors and Louvain clustering at 0.1 resolution. **c)** Percent representation of each cluster, stratified by donor group. “Low AD” signifies donors with an ADNC rank of No/Low, while “High AD” signifies donors with an ADNC rank of High. **d)** Dot plot representing expression levels of each marker gene by cell cluster in spatial transcriptomic dataset. **e)** UMAP of entire CosMx dataset following QC. **f)** Representation of each identified cluster, split between GM and WM. **g)** Single FOVs from spatial transcriptomics post-segmentation representing one WM (left) and one GM (right) FOV from the same donor. The white box signifies the enlarged inset region. **h)** OL cell segmentation with five ubiquitous OL transcripts mapped on the FOV. Each dot represents one *in situ* hybridization probe’s location within the tissue. **i)** The same OL segmentation with synapse- and neuron-related genes.

To contextualize WM transcriptomic findings within tissue architecture in different conditions, we performed spatial transcriptomics on Nanostring/Bruker’s CosMx platform using their 6000-probe set. We chose a subset of donors (n=6) from our WM cohort from whom multiple cortical regions have been evaluated in SEA-AD [31]. Like snRNA-seq, CosMx is an RNA-based analysis method, but it relies on *in situ* hybridization to label RNA transcripts on a thin section of tissue, preserving tissue architecture and providing spatial coordinates for each transcript and cell (**Supplemental Figure 1d**). For this study, we compared dorsolateral prefrontal cortex (DLPFC) samples from SEA-AD and WM samples performed specifically for this study. We selected high ADNC (n=3) and not/low ADNC (n=3) donors with DLPFC GM tissue. samples obtained adjacent to the region used for snRNA sequencing. Following data acquisition and QC to remove low-quality cells and fields of view, we identified each cluster’s cell type using candidate genes (**Fig. 1d-e)**. As expected, we found that OLs are the most prevalent WM cell type accounting for 87.6% of high ADNC and 85.1% of low ADNC cells, while GM was roughly a 1:1 neuron/glia split with 25.5% OLs in high ADNC and 24.6% in low ADNC GM (**Fig. 1f**). After each cell type was identified, we could label these cell types on representative fields of view, showing how broad cell type distribution varies between WM and GM areas (**Fig. 1g**). While WM was primarily composed of OLs, GM was more diverse, with multiple neuron subtypes and all glial types being represented. Because each *in situ* hybridization probe has spatial coordinates within the FOV, we mapped the distribution of five key myelination genes on the same WM and GM FOVs to visually correlate OL gene expression to transcriptomically identified OLs (**Fig. 1h**). Next, we assessed six key transcripts associated with the synapse or neurotransmitter cycling to highlight a key feature of GM. Surprisingly, we found GM OLs colocalized with these synapse transcripts more than WM OLs (**Fig. 1i**). Together, both snRNA sequencing and CosMx datasets identified similar WM cell type distribution, enabling specific downstream analyses employing both datasets.

### Integration of WM OLs and GM OLs

We combined our 70+ year-old donor WM OLs with the DLPFC GM OLs identified in SEA-AD (hereafter referred to as GM OLs for simplicity) to contrast the function of WM and GM OLs, highlighting regional variability in the OL population. We computationally isolated the GM OLs and integrated them with our WM OLs to form the UMAP in **Fig. 2a**. Next, we conducted differential expression (DE) and gene set enrichment analysis (GSEA) to identify unique features of WM and GM OLs, starting with myelination. We found comparable levels of myelin-associated gene expression (i.e. *MBP, MAG, MOG, MOBP, PLP1*) in WM and GM OLs (**Fig. 2b**), with significant enrichment of pathways involving autophagy and glial cell development in WM OLs relative to GM OLs (**Fig. 2c**).

**Figure 2.**
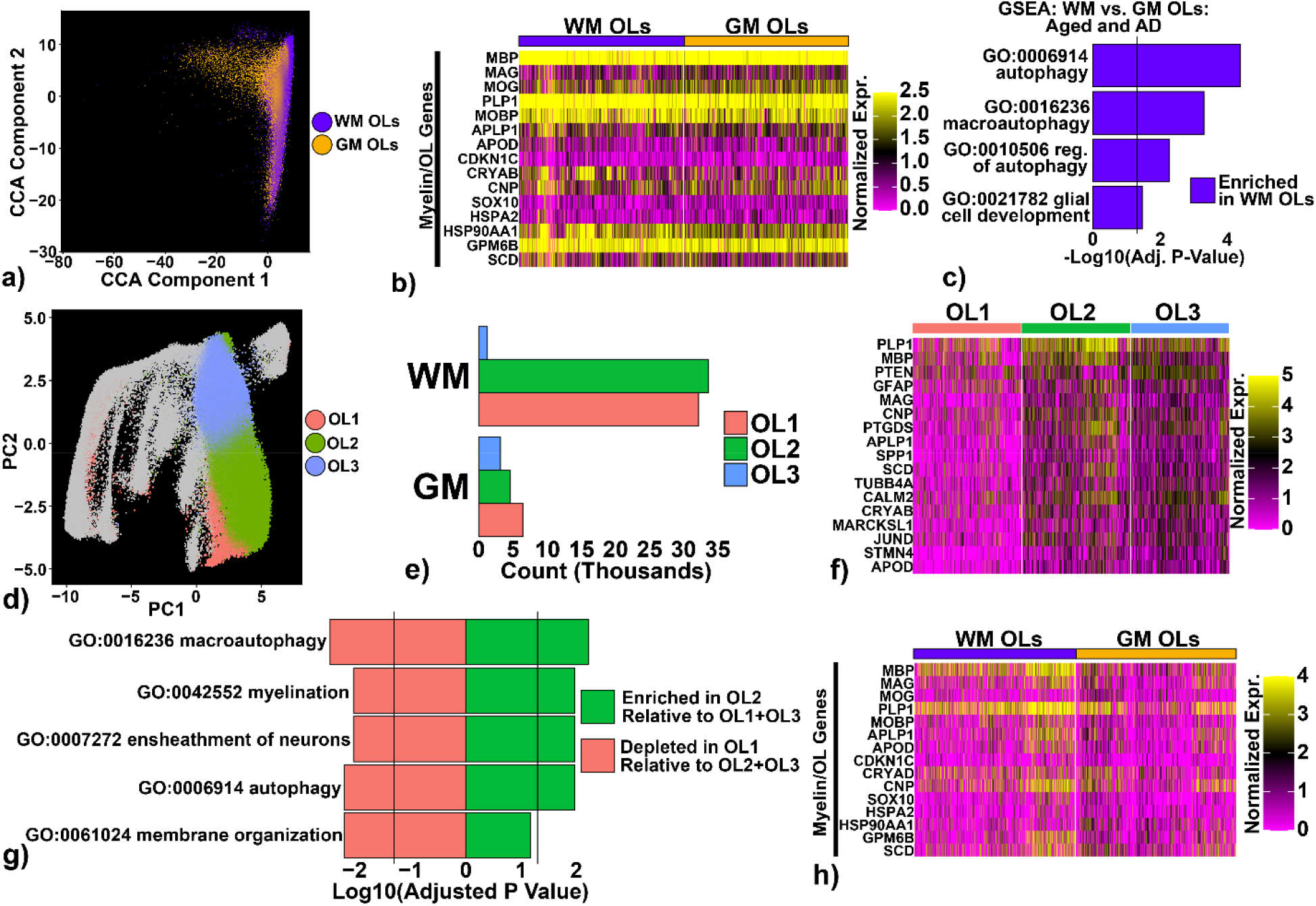
Myelin-related expression is similar between WM and GM OLs. **a)** UMAP following CCA-integration of 70+ year old WM and GM OLs. **b)** Heatmap showing expression of individual genes involved with myelination or OL identity. **c)** GSEA reveals significant enrichment of autophagy pathways in WM OLs relative to GM OLs. **d)** UMAP of CosMx dataset with OL clusters OL1, OL2, and OL3 identified. **e)** Distribution of OL1, OL2, and OL3 by tissue of origin. **f)** Expression of myelin and OL genes in OL1-OL3. **g)** GSEA reveals OL2 is significantly enriched for myelination and autophagy pathways, while OL1 is significantly depleted in these same pathways. **h)** Expression of myelin genes in WM and GM OLs in CosMx. Heatmap data is colored by normalized expression values of each gene, with each vertical column representing the expression within a single cell. GSEA results are reported as -Log_10_(adjusted p-values) following Benjamini-Hochberg correction. Vertical lines in GSEA plots represent an adjusted p-value of *q=0.05*.

Because we designed our CosMx experiments to assess similar WM and GM regions to our snRNAseq samples in a subset of the shared donors, we ran similar analyses on CosMx OLs. We found three unique OL clusters (**Fig. 2d**), with WM being primarily composed of OL1 and OL2 and GM containing a more even distribution of all three OL clusters (**Fig. 2e**). The expression of myelin-related transcripts was significantly different between OL clusters: OL2 showed upregulation (**Fig. 2f**) and enrichment (**Fig. 2g**) of these myelin transcripts, whereas OL1 showed downregulation and depletion. When OLs were divided by WM versus GM origin, we see a qualitative elevation of surveyed myelination genes in a subset of WM OLs, but this did not result in significant GSEA findings (**Fig. 2h**). Hence, our snRNA sequencing and CosMx data mutually support comparable levels of myelination transcript expression between GM and WM OLs.

### Synaptic Machinery in Human OLs

Recent studies have found evidence of OPCs directly modulating synapse structure and function [76], but the role of the OLs in these processes have not been clearly elucidated. While we showed that WM OLs express primarily myelination machinery, we hypothesized that GM OLs may be more involved with synapse and neurotransmitter functioning. During initial CosMx dataset characterization, we observed more synapse-related probes colocalized with GM oligodendrocytes (OLs) than with WM OLs (**Fig. 1i**). Consistent with this, GSEA in the snRNA-seq dataset showed that aged GM OLs were enriched for multiple synapse-associated processes relative to aged WM OLs, including regulation of trans-synaptic signaling, synapse organization, neurotransmitter transport, glutamatergic synaptic transmission, and the synaptic vesicle cycle (**Fig. 3a**). At the gene level, GM OLs showed broadly higher normalized expression of canonical synaptic/neurotransmission-associated transcripts compared with WM OLs in both snRNA-seq (**Fig. 3b**) and CosMx (**Fig. 3c**), supporting a reproducible regional difference across modalities.

**Figure 3.**
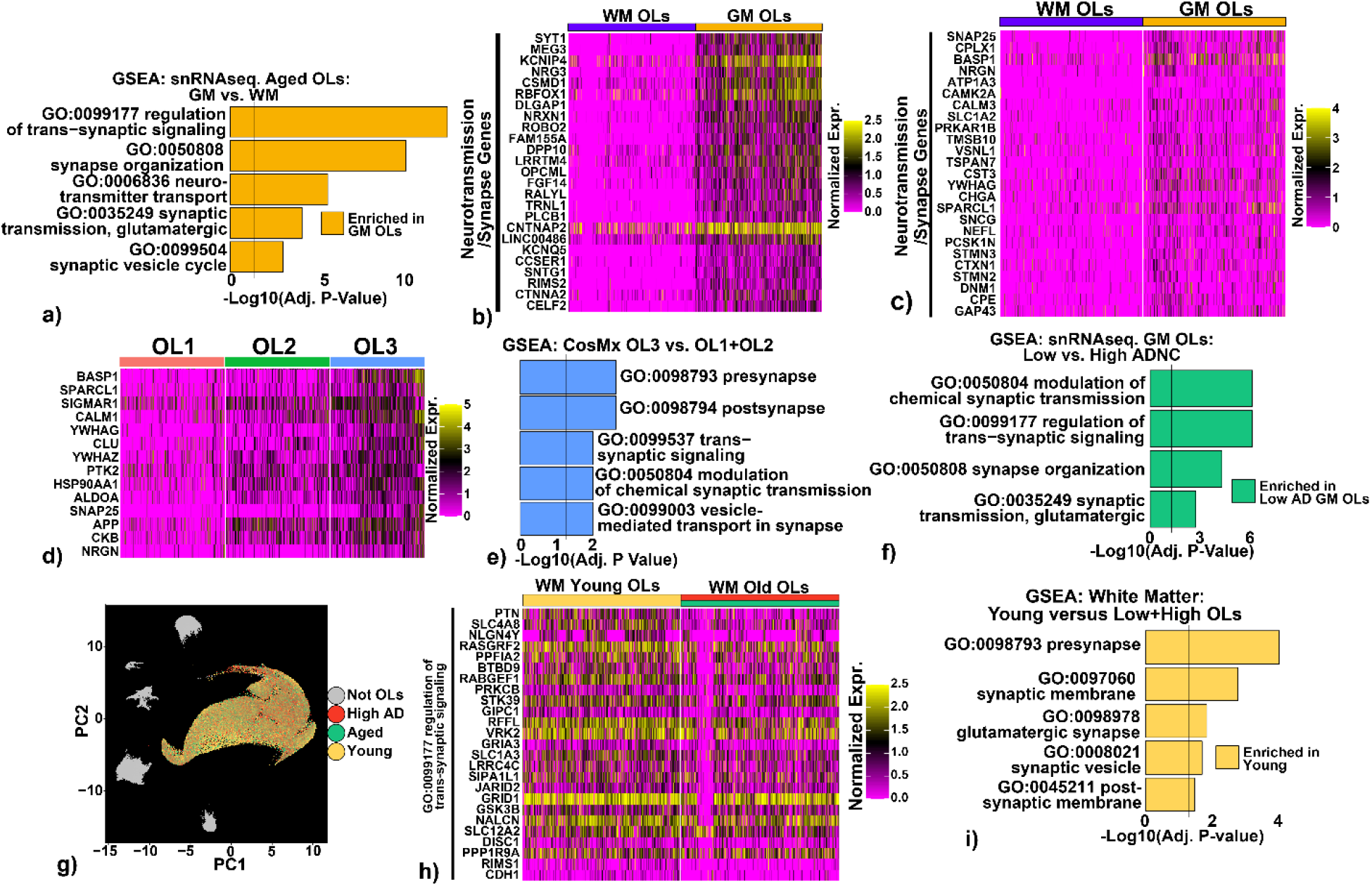
GM OLs show significant enrichment of synapse and neurotransmission-related transcripts. **a)** Aged (not/low and high AD pathology combined) GM OLs show enrichment of synaptic pathways relative to aged WM OLs in snRNAseq. **b-c)** GM OLs show upregulation of individual genes involved in neurotransmission and synapse function (right half of heatmap) relative to WM OLs (left half of heatmap) in both snRNAseq. **(b)** and CosMx **(c). d)** OL3 shows the most expression of these transcripts compared to the other clusters. **e)** GSEA analysis supports the expression pattern in 3d, finding enrichment of synapse machinery in OL3 relative to OL1 and OL2. **f)** Within old GM OLs, there was enrichment for synaptic pathways in no/low ADNC relative to high ADNC stage donors in snRNAseq. **g)** WM OLs are highlighted on the UMAP of the WM dataset. **h)** Young WM OLs contain more transcripts from genes included in “GO:0099177 regulation of trans-synaptic signaling” relative to old WM OLs. **i)** Young WM OLs show enrichment of synaptic pathways in GSEA analyses relative to old WM OLs (both aged and high AD OLs). Heatmap data is colored by normalized expression values of each gene, with each vertical column representing the expression within a single cell. GSEA results are reported as - Log_10_(adjusted p-values) following Benjamini-Hochberg correction. Vertical lines in GSEA plots represent an adjusted p-value of *q=0.05*.

Importantly, the CosMx OL compartment was not uniform: the OL3 cluster—predominantly GM-derived—showed the clearest synapse-associated expression pattern (**Fig. 3d**) and was specifically enriched for presynapse, postsynapse, trans-synaptic signaling, and vesicle-mediated transport in synapse gene sets relative to OL1+OL2 (**Fig. 3e**). This cluster-restricted enrichment supports the interpretation that synapse-associated OL transcriptional programs are concentrated in a discrete GM-biased OL subpopulation rather than representing a diffuse, uniform signal.

We next asked whether disease and age modulate this OL synapse-associated signature. In GM, synaptic gene sets such as chemical synaptic transmission, regulation of trans-synaptic signaling, synapse organization, and glutamatergic synaptic transmission were significantly enriched in low ADNC compared to high ADNC GM OLs (**Fig. 3f**), indicating attenuation of this program in high ADNC. In WM, young donors exhibited stronger synaptic signatures than older donors: young and aged/high-ADNC groups showed separation in a low-dimensional projection (**Fig. 3g**), and synapse-associated transcripts were generally higher in young WM OLs than in old WM OLs (**Fig. 3h**), with concordant enrichment of presynapse/synaptic membrane–related pathways in young versus combined aged WM OLs (**Fig. 3i**). Together, these results support a model in which GM OLs, particularly a GM-biased OL subpopulation, exhibit a prominent synapse-associated transcriptional program that diminishes with both aging and increasing AD neuropathology, consistent with reduced OL–neuron synaptic coupling or synapse-proximal OL states in vulnerable contexts.

### Immune Functions of OLs

Brain aging is characterized by a shift toward a more pro-inflammatory environment and altered immune signaling is a central component of the senescence-associated secretory phenotype (SASP) [22, 26, 40]. As such, we asked whether OLs exhibit measurable immune-associated transcriptional programs and whether these programs preferentially emerge with age and/or AD neuropathology. In WM snRNA-seq, GSEA comparing young versus aged WM OLs (no/low and high ADNC combined) revealed enrichment of immune-related pathways in aged OLs, including the HLA protein complex and multiple cytokine-response terms (**Fig. 4a**). Consistent with these pathway-level results, genes belonging to the “response to cytokine” Gene Ontology program showed broadly higher normalized expression in old WM OLs relative to young WM OLs (**Fig. 4b**), supporting a coherent age-associated immune-response signature within the WM OL compartment.

**Figure 4.**
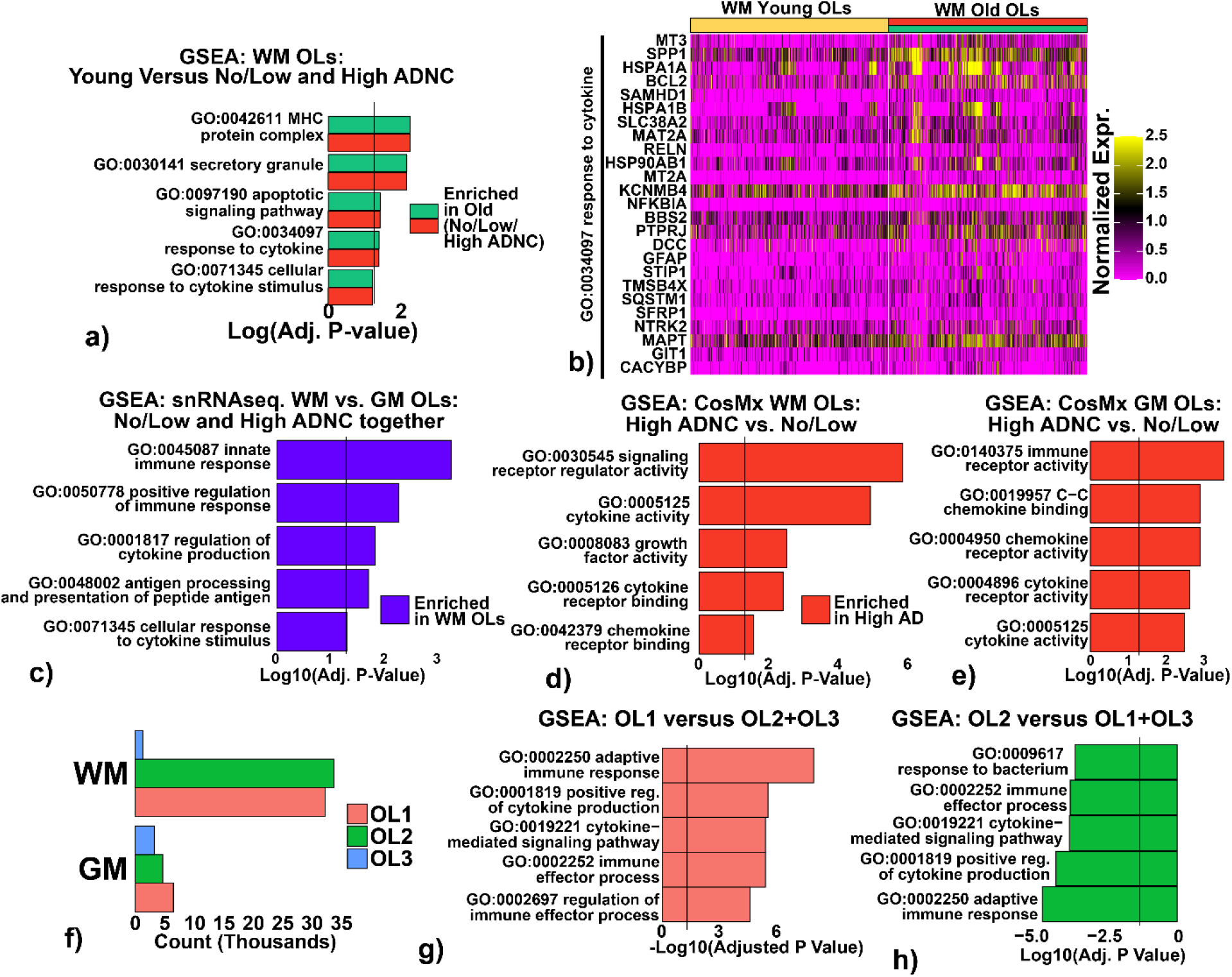
Human OLs are involved in immune processes, especially in the aged and AD brain. **a)** GSEA reveals aged and AD OLs show significant enrichment of MHC complexes and cytokine activity relative to young OLs in WM. **b)** Individual cell expression of genes included in the gene ontology pathway “GO:0035097 response to cytokine) are upregulated in a portion of the old WM OLs. **c)** WM OLs show significant enrichment of innate and adaptive immunity processes compared to GM OLs in snRNAseq. **d-e)** Following stratification of CosMx OLs by WM or GM origin, GSEA reveals that high ADNC donor OLs show enrichment for cytokine and chemokine pathways relative to not/low ADNC donor OLs in both WM **(d)** and GM **(e)**. **f)** significant enrichment for pathways involving adaptive immunity and cytokine activity. **h)** CosMx OL2 shows depletion of immune pathways involving cytokines and the adaptive immune response. Heatmap data is colored by normalized expression values of each gene, with each vertical column representing the expression within a single cell. GSEA results are reported as -Log_10_(adjusted p-values) following Benjamini-Hochberg correction. Vertical lines in GSEA plots represent an adjusted p-value of *q=0.05*.

Regional context further reinforced this pattern. In the WM vs GM OL analysis (not/low and high ADNC together), WM OLs were enriched for both innate and adaptive immune processes compared with GM OLs, including innate immune response, positive regulation of immune response, regulation of cytokine production, antigen processing and presentation of peptide antigen, and cellular response to cytokine stimulus (**Fig. 4c**). Thus, beyond aging effects, WM OLs were associated with stronger immune-related transcriptional programs.

These findings were recapitulated in the spatial CosMx dataset, where high ADNC OLs in both WM and GM showed enrichment for cytokine/chemokine signaling modules compared with no/low ADNC OLs. In WM CosMx OLs, high ADNC was associated with enrichment of signaling receptor regulator activity, cytokine activity, cytokine receptor binding, growth factor activity, and chemokine receptor binding (**Fig. 4d**). In GM CosMx OLs, high ADNC similarly increased enrichment for immune receptor activity and chemokine/cytokine signaling terms, including C–C chemokine binding, chemokine receptor activity, cytokine receptor activity, and cytokine activity (**Fig. 4e**). Consistent with tissue-level effects, WM was dominated by OL1 and OL2, whereas GM contained a larger contribution of OL3 (**Fig. 4f**), providing a cellular backdrop for state-specific immune differences.

Finally, immune pathway enrichment differed across CosMx OL clusters. OL1 showed relative enrichment for adaptive immune response and cytokine-related processes compared with OL2+OL3 (**Fig. 4g**), whereas OL2 showed depletion of these same immune-associated pathways relative to OL1+OL3 (**Fig. 4h**). Together, across snRNA-seq and spatial profiling, OLs exhibit robust immune-associated transcriptional programs that (i) increase with aging in WM, (ii) are stronger in WM than GM, (iii) are further enhanced in high ADNC in both tissues, and (iv) are not uniform across OL states, with immune enrichment concentrated in specific OL clusters.

### Evidence of Elevated Cell Stress in WM OLs from High ADNC Donors

The cellular senescence phenotype includes accumulation of DNA damage [38, 68, 77], chromatin remodeling [64, 68], lysosomal dysfunction [84], protein misfolding/aggregation [38, 65, 74], iron dysregulation [2, 57], and metabolic stress [43, 63], motivating a focused evaluation of proteostasis and damage-response programs in OLs across regions and ADNC burden. Stratifying aged OLs by anatomy and ADNC level revealed a clear WM-selective amplification of stress-protective transcripts in high ADNC, reflected primarily as higher expression magnitude per cell rather than a global increase in the fraction of expressing cells (**Fig. 5a**). WM OLs from high ADNC donors showed elevated scaled expression across multiple chaperone/proteostasis-associated genes (e.g., HSPA1A, HSP90AA1, DNAJB-family members, HSPH1, CHORDC1) alongside stress-linked factors such as TXNIP and iron/metal-handling genes (FTH1/FTL; SLC11A2) (**Fig. 5a**). When restricted to WM OLs, these patterns remained evident across the ADNC axis, with high ADNC exhibiting the strongest shifts relative to both young and not/low ADNC groups (**Fig. 5b**). These data were independently supported by violin plots highlighting increased expression of representative stress and iron-homeostasis transcripts (HSPA1A, HSP90AA1, FTH1, FTL, SLC38A2, CRYAB) (**Fig. 5c**). Consistent with these gene-level patterns, GSEA comparing high versus not/low ADNC identified enrichment of protein folding/chaperone and unfolded-protein response terms in both WM and GM (**Fig. 5d–e**), but heat shock protein binding was uniquely enriched in WM (**Fig. 5d**; *red box*), suggesting a more prominent HSR-like proteostasis arm in WM OLs under high AD pathology. Finally, WM versus GM comparisons in aged and AD OLs highlighted additional WM-skewed stress biology, including cellular response to starvation, double-strand break repair/DNA recombination (including homologous recombination), and mitochondrial autophagy (**Fig. 5f**). Collectively, these data implicate a model in which high ADNC is associated with heightened proteostasis demand and damage-adaptation programs in WM oligodendrocytes.

**Figure 5.**
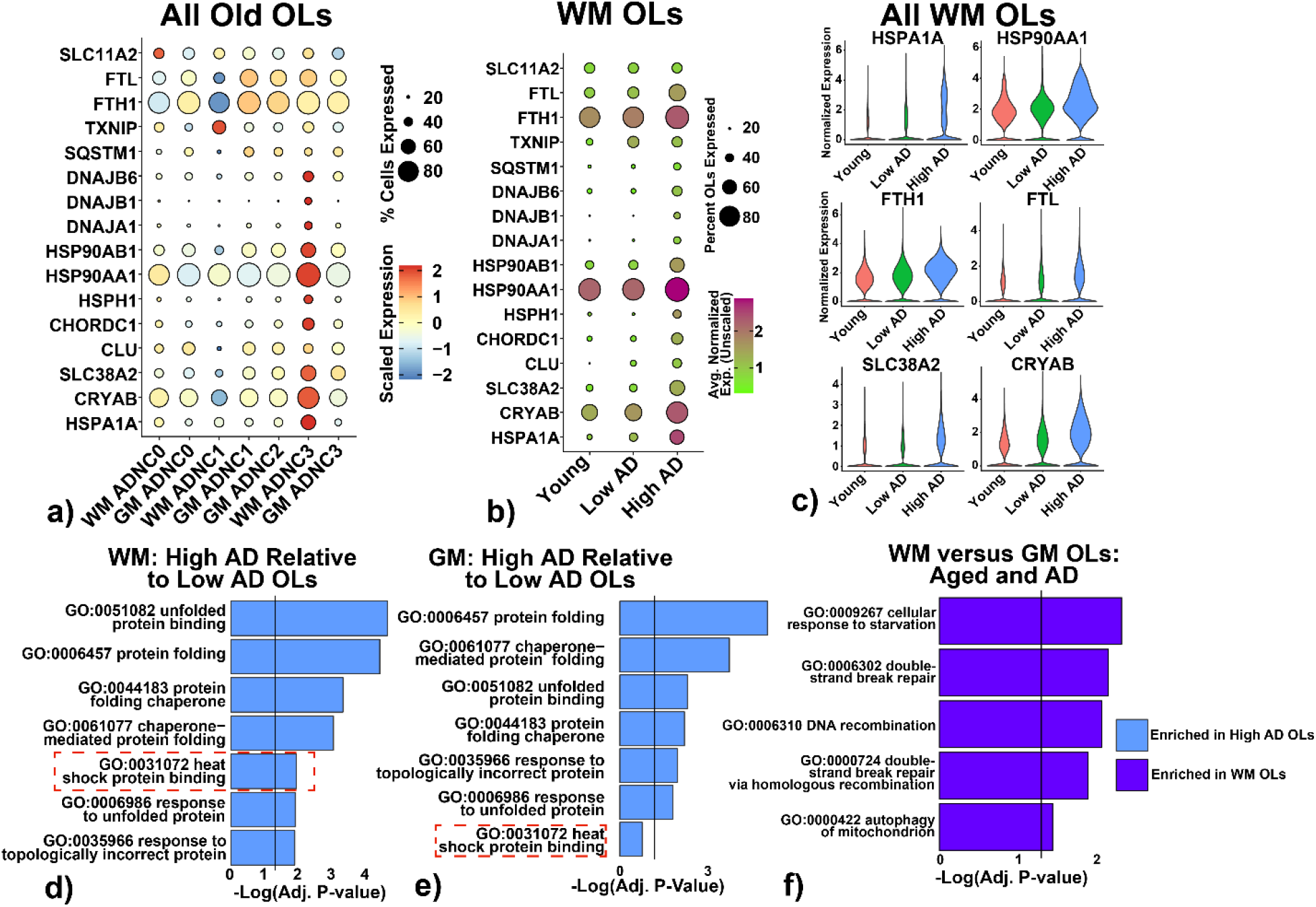
AD OLs show evidence of ER stress and misfolded proteins in both WM and GM relative to age-matched controls. **a)** Dot plot showing expression of key genes involved in un-/mis-folded protein response and heat shock proteins in WM high AD OLs. OLs were stratified by GM or WM origin, then by ADNC rank (0=Not, 1=Low, 2=Intermediate, 3=High AD pathology). Circle size indicates percentage of that cluster expressing each gene, while color represents relative expression to all other groups. WM ADNC=3 OLs show elevated expression of most of these heat shock proteins. **b)** Dot plot showing same HSPs across young, low AD old, and high AD old OLs. While young and low AD OLs show similar expression levels of these chaperone proteins, more high AD OLs express these transcripts and at higher levels. **c)** Violin plots of six of the most elevated transcripts in high AD OLs. All violin plots are showing only WM OLs stratified by young, low AD, and high AD groups. **d-e)** GO pathway enrichment in high AD WM OLs relative to young WM OLs **(d)** and low AD WM OLs **(e).** Both show enrichment of unfolded protein response and heat shock protein pathways in high AD WM OLs relative to young and low AD WM OLs. **f)** GO pathway enrichment in high AD GM OLs (ADNC=3) versus not/low AD GM OLs (ADNC=0-1). ADNC=2 GM OLs were excluded from this analysis. While high AD GM OLs show enrichment of the same unfolded protein response pathways as high AD WM OLs (blue bars), they do not show enrichment of heat shock proteins specifically (red dashed box), hinting at distinct un-/mis-folded protein responses in WM versus GM OLs. GSEA results are reported as - Log_10_(adjusted p-values) following Benjamini-Hochberg correction. Vertical lines in GSEA plots represent an adjusted p-value of *q=0.05*.

### Stress/ISR-reactive State Trajectories in High ADNC WM OL

Next, we annotated our unsupervised OL clusters using curated marker panels capturing (i) lineage stage, (ii) dominant stress/immune programs, (iii) lipid/cholesterol handling, (iv) proliferation, and (v) potential non-OL contamination to interpret OL state structure prior to pseudotime and directional-flow modeling. Canonical mature-myelin genes (e.g., *PLP1, MBP, MOG, MOBP*) identified terminal myelinating OLs, while an OPC/precursor panel (*PDGFRA, CSPG4/NG2, PTPRZ1*) was used only as an exclusion step to confirm minimal precursor contamination and to restrict downstream state modeling to OLs[29]. We defined a newly formed/differentiating OL (NFOL) panel enriched for *BCAS1* and *ENPP6*, supported by prior work describing *BCAS1* as a marker of early myelinating OLs and *ENPP6* enrichment in newly formed/differentiating OL populations[54]. We annotated a Stress/ISR-reactive program using canonical proteostasis/ER-stress and stress-response transcripts (e.g., *ATF3, HSPA1A/B, DNAJB1, DDIT3*), consistent with an integrated-stress/proteostasis response program in oligodendroglia[32]. We also included a lipid/membrane remodeling panel to capture shifts toward lipoprotein-associated lipid handling and membrane lipid remodeling reported in human oligodendroglia (*APOE, APOD, ABCA2, SCD, SGMS1, AMACR*)[45]. Based on dominant panel signals, we renamed clusters for downstream analyses as: Oligo_1 = NFOL/differentiating (*BCAS1/ENPP6+*), Oligo_2 = lipid remodeling (*APOE/ABCA2/LPL*), Oligo_3 = mature myelinating (MBP, MOG, MOBP), and Oligo_4 = Stress/ISR-reactive OLs (*ATF3, HSPA1A/B, DNAJB1, DDIT3*) (**Fig. 6a–b**). Because the mature myelinating population was much smaller than the other states, it was retained in the embedding visualization (**Fig. 6a–b**) but excluded from the subsequent state-to-state transition graphs to avoid unstable edge estimates dominated by low counts.

**Figure 6.**
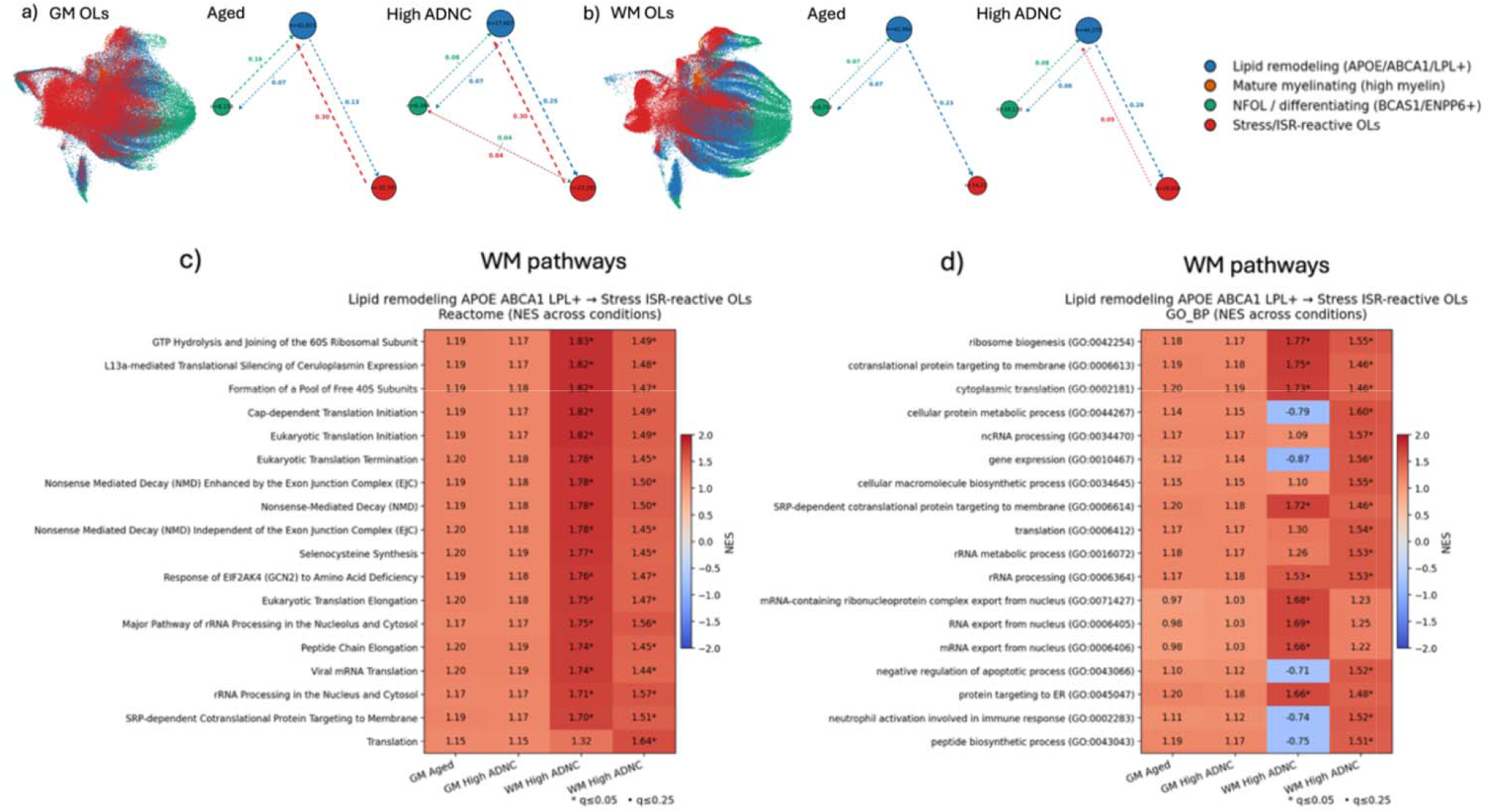
Oligodendrocyte-state structure and directional flows highlight a lipid-remodeling intermediate en route to ISR-reactive stress programs across tissue and AD strata. **(a) GM OLs:** Low-dimensional embedding of gray-matter oligodendrocytes colored by the four curated programs (NFOL/differentiating [BCAS1/ENPP6+], lipid remodeling [APOE/ABCA1/LPL+], Stress/ISR-reactive OLs, and mature myelinating [high myelin]). At right, directed state-transition graph inferred from within-stratum pseudotime (rooted in NFOL/differentiating) and kNN-based forward-edge “voting” (STRINGENT Δt ≥ 0.03), shown separately for GM No AD and GM AD; edge labels indicate normalized transition weights (fraction of forward edges from source state landing in the target state), and node labels indicate cells per state and edge colors denote the source state. **(b) WM OLs:** As in (a), but for white-matter oligodendrocytes, with directed transition graphs shown for WM No AD and WM AD. Across tissues, NFOL/differentiating is positioned as a root-like compartment coupled with the lipid-remodeling program, and lipid remodeling exhibits dominant forward flow into the Stress/ISR-reactive state; dotted arrows denote graph-supported multi-step flow when the dominant forward path is inferred along connectivity rather than a single strong direct edge. **(c) Reactome enrichment for lipid→ISR conversion:** Gene set enrichment analysis (GSEA) of transition-associated programs for the **lipid remodeling (APOE/ABCA1/LPL+)** → **Stress/ISR-reactive OLs** direction, showing Reactome pathways (normalized enrichment score, NES) across GM No AD, GM AD, WM No AD, and WM AD. **(d) GO Biological Process enrichment for lipid**→**ISR conversion:** As in (c), but showing GO BP terms (NES) across the same four strata. For enrichment panels, asterisks denote FDR q ≤ 0.05 and dots denote q ≤ 0.25.

We then computed a continuous pseudotime trajectory separately within each tissue stratum and condition (GM vs WM; Aged vs High ADNC), rooting the trajectory in the NFOL/differentiating (*BCAS1/ENPP6+*) compartment to infer directional state-to-state flows within the oligodendrocyte lineage (**Fig. 6a–b**). Across strata, the inferred topology supported a core architecture in which NFOL/differentiating and lipid remodeling are coupled, and the dominant forward flow from lipid remodeling proceeds toward a Stress/ISR-reactive state (**Fig. 6a–b**). In WM, the lipid→stress edge was prominent in both groups (0.23 in Aged; 0.20 in High ADNC) and coincided with a marked expansion of Stress/ISR-reactive OLs in High ADNC (n≈29,050) relative to Aged (n≈14,223), consistent with increased engagement of stress-response programs in diseased WM.

The reverse stress→lipid edge was weak/undetectable in Aged WM but became measurable in High ADNC WM (0.05), suggesting limited re-entry toward lipid remodeling under high pathology. In GM, stress↔lipid coupling was comparatively dynamic: the lipid→stress increased from 0.13 (Aged) to 0.25 (High ADNC). In addition, High ADNC GM showed low-weight direct connections between NFOL and Stress/ISR (0.04), consistent with additional (but minor) routes into/out of the stress program (Fig. 6a). Together, these patterns indicate that WM exhibits a pathology-linked expansion of the Stress/ISR compartment with only limited backflow, whereas GM maintains stronger bidirectional cycling between lipid remodeling and Stress/ISR across conditions (Fig. 6a–b).

Finally, pathway enrichment of transition-associated gene programs positioned the *APOE/ABCA1/LPL*+ lipid-remodeling state as an intermediate en route to Stress/ISR-reactivity, with enrichment summarized for the lipid remodeling → Stress/ISR-reactive transition (**Fig. 6c–d**). Across strata, Reactome terms were dominated by translation and ribosome-associated programs such as eukaryotic translation initiation/ elongation/ termination, ribosome subunit joining, nonsense-mediated decay, rRNA processing, and SRP-dependent cotranslational targeting, while GO Biological Process terms highlighted ribosome biogenesis, cytoplasmic translation, ncRNA/rRNA processing, RNA export, and protein targeting to ER (**Fig. 6d**). Notably, several broad biosynthetic/RNA-metabolism terms showed a WM-specific shift with pathology in the form of depletion in WM Aged but enrichment in WM High ADNC in Fig. 6d, consistent with ADNC linked rewiring of proteostasis and RNA-handling demands during the lipid→stress/ISR conversion. Altogether, the inferred transitions suggest that WM OLs in high ADNC are preferentially routed toward a Stress/ISR-reactive state with limited reverse flow, whereas GM retains stronger bidirectional coupling between lipid remodeling and Stress/ISR programs (**Fig. 6a–b**). The accompanying enrichment profiles indicate that entry into Stress/ISR-reactivity is accompanied by increased translation/ribosome and RNA-metabolism programs, consistent with elevated biosynthetic and proteostasis burden during this state conversion (**Fig. 6c–d**). A model that incorporates the findings of this study is shown in **Figure 7**.

**Figure 7.**
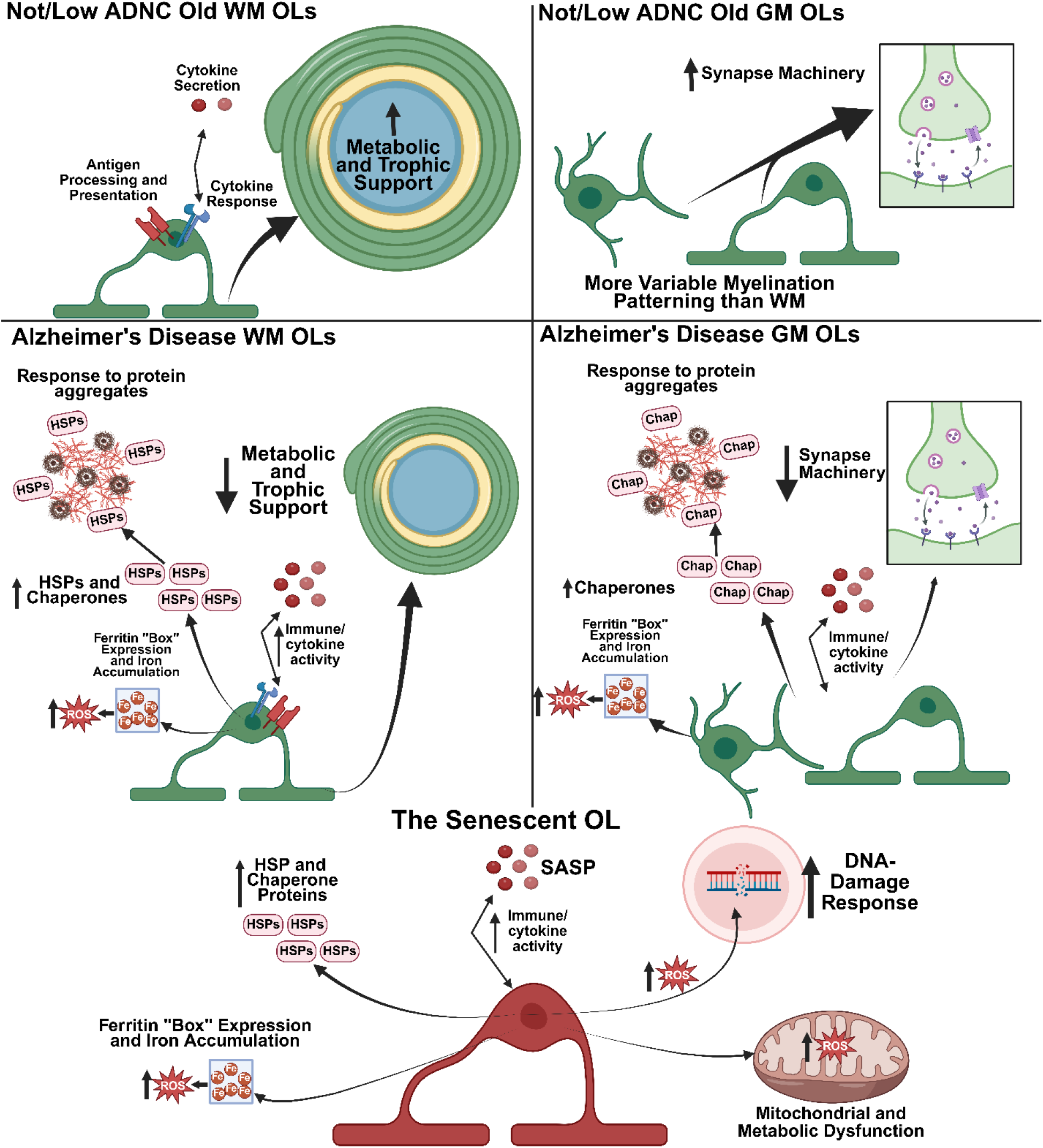
Schematic summarizing the findings from the snRNAseq. and CosMx data. The top half of this cartoon depicts the identified roles of WM OLs (left side) and GM OLs (right side) in the context of not/low ADNC pathology (top panels) and high ADNC pathology (bottom panels). The bottom portion of this cartoon summarizes the proposed phenotype of the senescent OL, potentially present in both WM and GM. This figure was made in BioRender.

## Discussion

Although white matter (WM) comprises roughly half of the human brain, its cellular composition and disease-relevant biology remain less well-resolved than those of grey matter (GM)[89]. Here, we generated a WM-centric single-nucleus RNA-seq dataset complemented by spatial transcriptomics (CosMx) with a specific focus on oligodendrocytes (OLs), enabling direct comparison across anatomical region (WM vs GM), age, and AD neuropathologic change (ADNC). Consistent with the expected cellular architecture of WM, myelinating OLs comprised the majority of captured WM cells (**Fig. 1**), providing a robust substrate for OL-resolved analyses across modalities.

A central observation from WM-GM integration was that core myelin transcript abundance did not simply scale with regional myelin content. Despite the well-established structural differences between WM and GM myelination [85, 87], WM and GM OLs showed broadly comparable expression of canonical myelin genes (**Fig. 2**), while WM OLs exhibited distinct pathway-level differences, including enrichment for autophagy and glial-development programs (**Fig. 2**). Together, these findings suggest that steady-state transcription of core myelin genes is relatively similar across regions, whereas WM OLs differ more prominently at the level of homeostatic maintenance and stress-handling programs than at baseline myelin transcript levels. Beyond myelination, our data support a GM-biased OL synapse-associated transcriptional program that is reproducible across snRNA-seq and CosMx (**Fig. 3**).

GSEA and gene-level visualization showed that aged GM OLs are enriched for synapse-associated processes relative to aged WM OLs (e.g., regulation of trans-synaptic signaling, synapse organization, neurotransmitter transport, glutamatergic synaptic transmission, and synaptic vesicle cycling; **Fig. 3a–c**). Importantly, this signal was not uniform within the OL compartment: a GM-enriched CosMx OL subcluster (OL3) carried the strongest synapse-associated expression pattern and pathway enrichment relative to other OL clusters (**Fig. 3d–e**), indicating that this signature is concentrated in a discrete OL subpopulation rather than diffusely expressed across the lineage. This program was attenuated with higher ADNC in GM (**Fig. 3f**) and was stronger in young WM OLs than in aged WM OLs (**Fig. 3g–i**), consistent with the possibility that OL states associated with synapse-proximal functions (or activity-responsive programs) are reduced with aging and AD pathology. These observations align with prior evidence that OL-lineage cells can sense neuronal activity and engage ion-channel and synapse-adjacent machinery to modulate myelin remodeling [8, 41, 46] while leaving open key mechanistic questions about whether OL synapse-associated transcripts reflect functional synapses, synapse-proximal localization, or broader activity-responsive states. There is evidence that OLs express Kir4.1 channels (product of *KCNJ10* gene) that help regulate extracellular potassium levels at the site of nodes of Ranvier and that inflammatory states can negatively alter this mechanism [47]. It is plausible that the pro-inflammatory environment of the aged and AD OLs is causing similar disruptions as we found differential expression of other KCN genes that are involved with potassium channels.

In parallel, we observed robust evidence that OLs engage immune-associated transcriptional programs that vary with age, tissue context, disease, and OL state (**Fig. 4**). In WM snRNA-seq, aged OLs showed enrichment of immune-related pathways relative to young OLs, including MHC complexes and cytokine-response processes (**Fig. 4a–b**). When GM and WM OLs were analyzed together, WM origin was associated with stronger immune-related enrichment than GM (including innate immune response, regulation of cytokine production, antigen processing/presentation, and cellular response to cytokine stimulus; **Fig. 4c**). These trends extended to spatial data: high ADNC CosMx OLs in both WM and GM showed enrichment for cytokine/chemokine signaling modules relative to not/low ADNC (**Fig. 4d–e**), and immune enrichment was state-dependent, with OL1 showing relative enrichment for immune processes compared with other OL clusters while OL2 showed depletion (**Fig. 4g–h**). These findings suggest that aging and high ADNC expand immunologically engaged OL states, potentially reflecting either heightened responsiveness to an inflammatory parenchyma or a SASP-like phenotype restricted to specific OL states. There is strong evidence that the aging and AD brain becomes pro-inflammatory from a complex interplay of microglial activation and dysfunction [1, 42, 59, 70, 75], protein aggregation [53, 66], and astrocyte dysfunction [52, 62] (for comprehensive summary, please see [17, 20]). Notably, OL-derived lipid (myelin) contributes to microglia congestion and activation [42, 75], but our evidence suggests that OLs may be directly involved in this pro-inflammatory shift through cytokine secretion and activation based on our gene expression data (Figure 4). Further studies to elucidate whether cytokines are produced and released from OLs are warranted.

Multiple additional findings converge on a model of WM-selective amplification of proteostasis and damage-adaptation programs in AD (**Fig. 5**). High ADNC WM OLs displayed increased expression magnitude of a specific subset of stress-protective transcripts (including prominent chaperone/proteostasis genes and iron-handling factors such as ferritin subunits) (**Fig. 5a–c**). At the pathway level, both WM and GM OLs in high ADNC showed enrichment for protein folding/chaperone and unfolded-protein response terms (**Fig. 5d–e**), but WM uniquely showed enrichment for heat shock protein binding (**Fig. 5d**), suggesting a stronger or more specific heat-shock-like proteostasis response in association with high pathology. WM OLs also showed enrichment of pathways consistent with broader damage adaptation, including cellular starvation responses, double-strand break repair (including homologous recombination), and mitophagy (**Fig. 5f**). Prior studies have highlighted microvascular changes and iron-associated cell stress (ultimately leading to ferroptosis) in aging WM OLs [1, 6, 7], and our studies support these findings. A recent study found that cerebral amyloid angiopathy was associated with elevated intra-parechymal iron levels, and that these elevated iron levels correlated with faster cognitive decline [3]. It is plausible that accumulation of iron in OLs is responsible for these measured changes as OLs are the largest reservoir of iron in the brain [72].

These stress-response features overlap conceptually with components of proposed senescence-like programs, particularly when senescence is defined broadly to include maladaptive damage-response states in terminally differentiated cells [68], although they do not establish classical senescence. The combination of high metabolic demand [10, 12, 60, 79], large iron concentrations [1, 14, 18], and poor oxygen supply present in aged [25] (and further exacerbated in AD [83]) WM may provide a mechanism for why WM OLs express more damage adaptation and proteostasis machinery. When these stressors accumulate beyond the adaptive capabilities of the OL, they can result in autophagy and cell death (potentially through ferroptosis) [1, 6, 7, 9, 15]. Our data suggest that WM OLs are expressing more autophagy machinery which may indicate increase myelin turnover as well (Fig. 2 c, g).

To better resolve how these stress- and lipid-associated programs relate to OL state structure, we annotated OL clusters using literature-supported marker panels and applied pseudotime-based directional-flow modeling (**Fig. 6**). This analysis supported a consistent NFOL/differentiating → lipid remodeling → stress/ISR-reactive architecture, with high ADNC in WM marked by expansion of the stress/ISR-reactive compartment and strong lipid→stress directionality (**Fig. 6a–b**). Enrichment of transition-associated programs for the lipid remodeling → stress/ISR-reactive conversion emphasized translation/ribosome and RNA-handling pathways (**Fig. 6c–d**), consistent with increased biosynthetic and proteostasis demand during this conversion. Together with the WM-selective proteostasis, DDR, and mitophagy enrichments (**Fig. 5**), these results support a model in which high ADNC biases WM OLs toward persistent stress-adaptation states rather than transient responses.

A unifying theme of our findings is the intersection of oxidative stress, proteostasis strain, iron handling, and inflammatory signaling as reinforcing loops contributing to OL dysfunction. OLs have unusually high metabolic demands and iron requirements, and iron accumulation can potentiate oxidative damage and stress-response activation[1, 4–7, 10, 12–14]. Sustained activation of damage-response programs (DDR/UPR/proteostasis pathways) can, in turn, promote inflammatory signaling and cytokine production[33, 74, 88], potentially linking internal OL stress to parenchymal inflammation and SASP-like outputs. Although causality cannot be inferred from post-mortem tissue, the consistency across modalities and state-resolved analyses provides a coherent framework for future studies to test whether these programs represent compensatory adaptation, maladaptive persistence, or a senescence-like terminal trajectory in WM OLs. We summarize this working model contrasting homeostatic OL functions with aging/high-ADNC shifts toward stress/immune activation, iron/ROS burden, and impaired support functions in **Fig. 7**. While our present work was limited in scope to Oligodendrocytes, future work will expand our analysis to additional cell types to explore cellular composition and interactions in WM aging and disease.

Across complementary snRNA-seq and spatial CosMx profiling, our analyses define a WM-centered OL aging/ADNC program that extends beyond baseline differences in myelin transcripts. GM OLs, particularly a GM-biased OL subpopulation, exhibited a prominent synapse-associated transcriptional program that diminishes with higher ADNC, while immune-associated and stress-adaptation programs increase with aging and pathology and are concentrated in specific OL states rather than uniformly distributed across the lineage (**Figs. 3–6**). High ADNC further amplified WM-selective proteostasis, iron-handling, DNA-repair, starvation, and mitophagy signatures (**Fig. 5**) and coincided with expansion of a Stress/ISR-reactive compartment within a reproducible NFOL→lipid remodeling→Stress/ISR state architecture (**Fig. 6**). Together, these results support a model in which WM OLs are preferentially driven into persistent stress- and immune-associated states as a response to aging and/or high ADNC; features consistent with senescence-linked dysfunction and providing a mechanistic framework for targeted validation and therapeutic interrogation (**Fig. 7**).

## Supporting information

Full donor list

## Author Contributions

JPV and CDK conceptualized the projects and hypotheses within the paper. Methodology including WM nuclei isolation, 10x Genomics procedure, NanoString, and analysis scripts were designed by JPV, JARB, SRK, AMW. SRK, BFK, CLR were responsible for sequencing. Software development (R and python scripts) were written (and troubleshooting supported by) JPV, JARB, CLR, BFK, SRK, JAM. Experiments and formal analysis of single cell data were conducted primarily by JPV with pseudotime analysis from JARB. Analysis was supported by JAM and SRK. Tissue samples (brain collection, acquisition of cores, preparation of slides, etc.) was a team effort by the BRaIN lab at the University of Washington and authors AMW, AK, LK, CDK, MD, DDN, AN, CSL. The manuscript was written by JPV and JARB with review and editing from JPV, JARB, SRK, AMW, CDK, CSL, AN, DDC, JAM, SAB, LSS. Figures were generated by JPV (1-5,7) and JARB (6). Supervision was provided by CDK, SRK, AMW, JAM. Funding for reagents and materials was acquired for this project by CDK with individual support as described in funding section for individual authors.

All authors have reviewed and agreed with the final submitted manuscript.

## Acknowledgements

We thank the King County, Snohomish County, and Pierce County Medical Examiners and their offices and staff without whom the Pacific Northwest Brain Donor Project would be impossible. We thank Lisa Keene, Emily Ragaglia, Aimee Schantz, and Kathryn Torrez for incredible administrative support, John Campos and Mark Montine for data management, and Jenna Kelley, Julia Ryan, Flavia Ernau, Kim Howard, Chase Smith, Ming Hu, Tejas Bajwa, and Katie Miller for outstanding technical support. We appreciate Christine Mac Donald, Swati Rane, Eric Larson, Jeff Iliff, John Neumaier, Jessica Young, Brian Kraemer, Emily Schneidereit, Vishal Nigam, Sumie Jayadev, and Katie Prater for helpful discussions. Finally, we are deeply grateful to the research participants and their families without whom this work would be impossible.

## Methods

### University of Washington BioRepository and Integrated Neuropathology (BRaIN) Laboratory

The UW BRaIN lab supports numerous prospective cohort studies of aging, dementia, and neurodegeneration including the UW Alzheimer’s Disease Research Center (ADRC), the Kaiser Permanente Washington Health Research Institute Adult Changes in Thought (ACT) study, and others. In addition, as part of our work with the BRAIN Initiative Cell Atlas Network, we leverage the Pacific Northwest Brain Donor Network to identify potential normal adult brain donors from age 16-70 through local medical examiner offices. Once identified, in coordination with the ME offices, potential donor families are approached to consider brain donation. All brain donors have provided informed consent and/or have the consent confirmed by legal next of kin after the donor has passed away. The BRaIN lab methods are described elsewhere [49]. Briefly, in late 2018 methods for procurement, dissection, preservation, and characterization were updated to include thin (4mm) fresh brain slicing, either rapid freezing in supercooled isopentane or fixation in 10% neutral buffered formalin in alternating slabs, a procedure we refer to as ‘precision rapid procurement’. Fixed brain slabs as sampled extensively following the “network-based autopsy” protocols developed in the BRaIN lab to capture not only classical neuropathology and comprehensive neurodegenerative disease neuropathology, but also neuroimaging and functionally relevant connectivity nodes and tracts within the human brain[49]. All donors in the BRaIN lab, including those aged 16-70, undergo the same comprehensive workup that includes extensive staining for ADNC, other tauopathies including chronic traumatic encephalopathy, TDP-43 pathology including limbic-predominant, age-related TDP-43 encephalopathy LATE), synucleinopathies including Lewy body disease, and extensive workup for microscopic and macroscopic vascular brain injury and vasculopathies.

### Aged Donor Selection – White Matter

Aged brain donors were selected from the UW BRaIN Lab using the following criteria: RNA Integrity Number (RIN) >7, fresh frozen tissue available using optimized methods (isopentane freezing implemented in late 2018), comprehensive neurodegenerative neuropathology workup (including ADNC score, Lewy body disease, limbic-predominant age-related TDP-43 encephalopathy (LATE) pathology, and a variety of vascular pathology/vascular brain injury metrics), and age >70. From this list a group of donors with high ADNC was identified and a demographically matched cohort of donors with not/low ADNC was identified. ADNC score was chosen as the primary AD metric because it is standard practice and incorporates Thal phase (amyloid plaque distribution), Braak score (neurofibrillary tangle distribution), and CERAD score (neuritic plaque density) into a composite ADNC score[44]. LATE stage 3 (TDP-43 detected in middle frontal gyrus)[61] donors were excluded, as were any brains with evidence of major cerebrovascular pathology like stroke or hemorrhage. Donors with Down syndrome, chronic traumatic encephalopathy, motor neuron disease including ALS, or multiple sclerosis were excluded as well. Most of the donors selected were from the Adult Changes in Thought (ACT) Study, a population-based longitudinal cohort that enrolls participants on a rolling basis, providing annual or biannual evaluations to assess cognitive status and medical history (https://www.actagingresearch.org/). Thus, the records we used were robust and most major medical history was assessed during selection. Following these criteria, a cohort of 32 donors was selected, with 19 (11 female, 8 male) with ADNC, and 13 (8 female, 5 male) with No/low ADNC. The summary of this cohort and their pathological scores are provided in **Supplemental Table 1**. All donor information is provided in **Supplemental Table 2**.

### Young Adult Cohort Case Selection – White Matter

The young adult cohort were derived from the Pacific Northwest Brain Donor Network and included 16 donors aged 50 years or under with minimal-to-no neuropathologic findings, including no ADNC. From the larger cohort of potential normal adult donors, initial inclusion criteria for this study included all donors aged 27-50 years with fresh frozen tissue available, RIN >7, postmortem interval <24 hours, and year of death in late 2018 or more recent to ensure precision rapid protocol was performed for tissue acquisition and preservation. From this list, donors were further selected to age and sex match as well as possible. Exclusion criteria for the young cohorts included schizophrenia and other serious psychiatric diagnosis, developmental disorders including autism, history of epilepsy, other significant neurological disorder, traumatic brain injury (if noted in records and/or requiring hospitalization), any concerning neuroinflammation, infection, or neoplasm, and any significant neurodegenerative disease or other neuropathological abnormalities, including evidence of acute/chronic hypoxic-ischemic injury, autolytic changes. While great care is taken to identify neurotypical brain donors, this is a very high bar so some level of age-related neuropathology and clinical diagnoses such as opioid dependence, chronic depression, etc. are tolerated. The summary statistics of this cohort are provided in **Supplemental Table 1**. All donor information is provided in **Supplemental Table 2**.

### Tissue Sampling

The general experimental schematic is displayed in **Supplemental Figure 1A**. All samples were procured from unfixed isopentane-frozen tissues kept on blocks of dry ice during sampling to ensure all tissues remained completely frozen throughout process. Samples were removed from -80°C freezer, placed on dry ice block, and cored with an 8-10mm sterile prechilled biopsy punch. The anatomic region targeted for all samples was roughly 0.5cm anterior to the anterior horn of the lateral ventricle, targeting the deep frontal white matter. Care was taken to ensure the entire core was white matter, not gray matter, and all samples were imaged after coring for record-keeping. These cores were bisected into two pieces, one piece for bulk RNA extraction and RIN calculation, the other for the nuclei isolation and 10x procedure (if RIN value was greater than 5.9). Note, RIN values above 5.9 were deemed sufficient for these experiments because all donors had previously been assessed for RNA integrity in both frontal and occipital poles as part of routine workflow, and thus this second RIN calculation was to ensure the sample itself was adequate to proceed. White matter RIN values were consistently 1-2 points below the gray matter RINs, thus we lowered our RIN cutoff to 5.9 accordingly for these WM samples.

### Nuclei Isolation

Nuclei isolation was conducted following the protocol developed by the Allen Institute and published through protocols.io (https://www.protocols.io/view/isolation-of-nuclei-from-human-or-nhp-brain-tissue-ewov149p7vr2/v3). However, since this protocol was designed for gray matter nuclei isolation, a few adjustments were necessary. First, myelin debris removal was done via an 21% iodixanol + nuclei suspension gradient laid over 25% iodixanol and spun at 8,000xg for 15 minutes at 4°C, rather than using myelin-binding beads. NeuN labeling and sorting was not performed, as our focus was OL, there were few neurons to begin with, and it was not desirable to alter the natural distribution of cell types. In brief, the sample was introduced to 1mL of RNAase inhibitor and protease inhibitor containing nuclei isolation media (NIM) and dounced using a sterile Eppendorf mortar and pestle on ice until all tissue was homogenized.

Homogenized samples were passed through 100um, 70um, and 30um filters, spun at 900xg for 10 minutes at 4°C, and supernatant removed. Samples were resuspended in equal parts NIM and 42% iodixanol to create a 21% iodixanol+nuclei solution, which was laid over ice-cold 25% iodixanol and spun at 8,000xg for 15 minutes at 4°C to isolate nuclei from myelin debris. The myelin debris/layer was removed with a sterile Q-Tip, and all supernatant removed via pipette. The resulting pure nuclei were washed 3 times in NIM to remove iodixanol and any residual myelin debris. An aliquot of the pure nuclei was stained with 1:1000 DAPI for 5 minutes and an aggregate of 3 automatic cell counts using the Countess II Cell Counter (Invitrogen: AMQAX1000) and 2 manual hematocytometer counts were averaged to obtain the nuclei concentration for each sample, along with purity (representative image of purified nuclei, Supplemental Figure 1B). These values were used for subsequent calculations regarding loading the 10x chip.

### 10x Procedure

The 10x Genomics procedure followed the Chromium Single Cell 3’ Reagent Kits User Guide (v3.1 Chemistry Dual Index) protocol, User Guide CG000315 (https://www.10xgenomics.com/support/single-cell-gene-expression/documentation/steps/library-prep/chromium-single-cell-3-reagent-kits-user-guide-v-3-1-chemistry-dual-index). During this process, each sample was loaded for a total cell capture of 8,000 cells per sample. Samples were run in groups of 2 or 4 per run, with samples within the run being samples from both the AD and Not/low AD groups to help account for batch effects. The young cohort was run separately from the old cohort, with the young non-fentanyl overdose and fentanyl overdose cases being included in all runs, again to help reduce batch effects due to individual run variability. PCR cycle number (Step 2.2d in protocol) was ran for 12 cycles, and the PCR cycle number (Step 3.5 in protocol) was always calculated based on quantified cDNA via Agilent Bioanalyzer High-Sensitivity DNA kit (PN: 506-4626) measurements for 200-9,000 base pair peaks (Sup. Figure 1C). Final cDNA library concentrations were determined using both Bioanalyzer High-Sensitivity DNA measurements (Sup. Figure 1C) and Invitrogen Qubit dsDNA Quantification Assay (PN:Q32854) measurements to determine both average base pair size and concentration of each cDNA sample library. The molarity of each cDNA library was calculated and samples were merged into sets of 4 samples of equimolar concentration to sequence. **Supplemental figure 1** provides representative images of tissue sampling, nuclei counts, and bioanalyzer results for a single run to demonstrate what a typical run looked like.

### Sequencing, Demultiplexing, Quality Control

Sequencing was done in sets of four samples using Illumina NovaSeq 6000 SP Reagent Kit v1.5 (100 cycle) (PN:20028401) sequencing kits. The raw sequencing data was run through 10x’s CellRanger (v7.2) pipeline to map to the Human reference genome HG38, demultiplex each sample, and provide initial quality control on samples. All cells with less than 200 features (genes) and greater than 5% mitochondrial reads were removed. Sequencing metrics like RIN values, cells captured, median genes per cell, and mean reads per cell are depicted in Supplemental Figure 3.

**Supplemental Figure 3.**
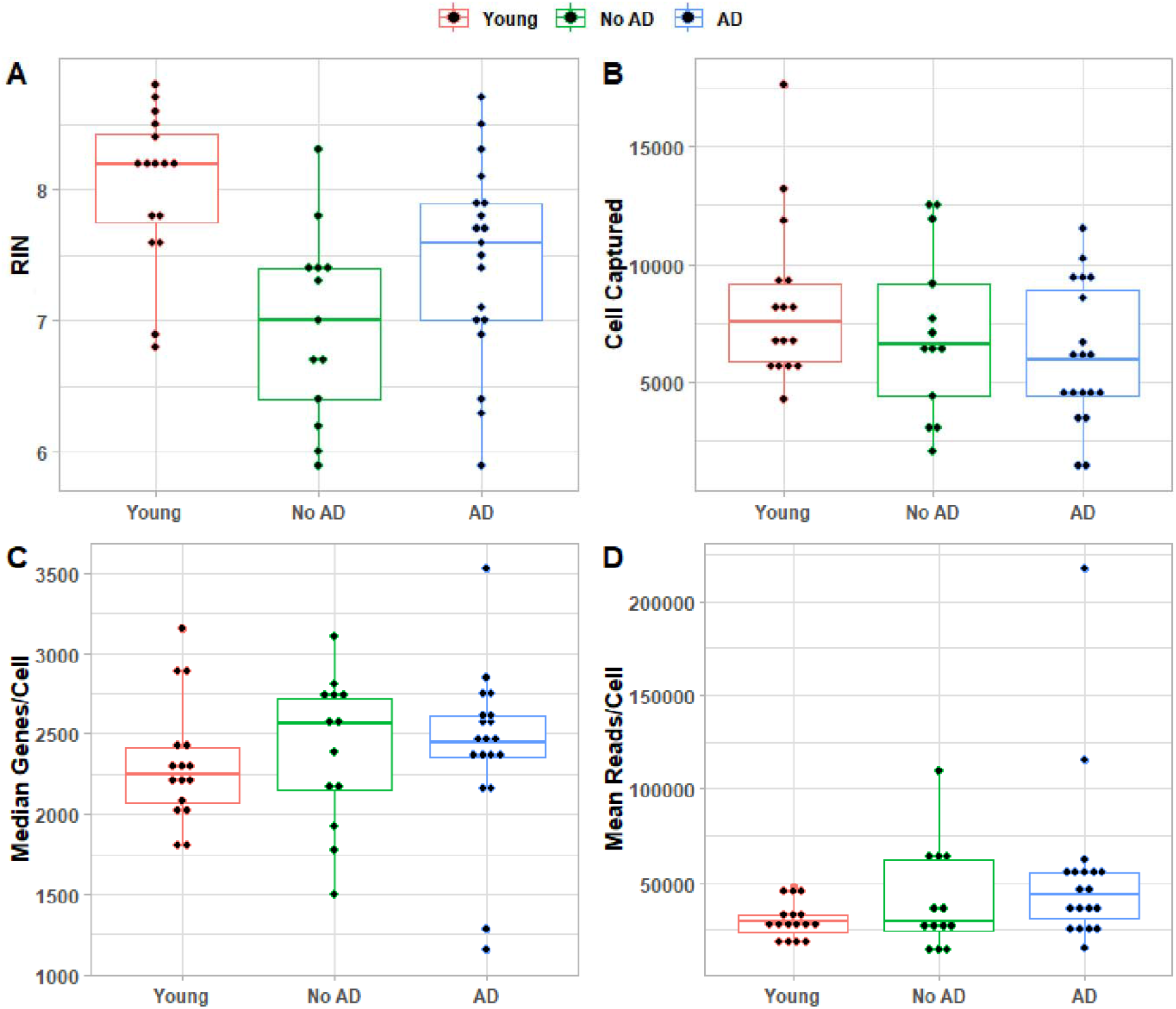
(QC metrics, if included as a supplemental figure) | Library quality and sequencing performance across donor groups. Boxplots summarizing sample- and library-level QC metrics across cohorts (Young, No AD/low ADNC, and high ADNC), including **(A)** RNA integrity number (RIN), **(B)** nuclei/cells captured per library, **(C)** median genes detected per cell, and **(D)** mean reads per cell. Each point represents an individual donor/library; boxplots summarize the distribution within each group and illustrate cohort-wide comparability of sequencing performance and inclusion thresholds used for downstream analyses.

All downstream analysis was conducted in RStudio using the Seurat (v5) package. All scripts are available on our GitHub (https://github.com/UW-BRaIN-lab). Individual HDF5 files taken from the CellRanger outputs were converted to RDS files, relevant metadata added, and the standard Seurat pipeline was run on each individual sample (normalize, FindVariableFeatures (nFeatures=2,000), scale, PCA, UMAP (dimensions 1:50, resolution=2 for pre-integrated clusters, resolution =1 for post-CCA integrated clustering). A note on this process: the normalization was done on each sample, rather than the whole dataset because this information was needed to run the DoubletFinder() process. DoubletFinder() is an algorithm that estimates the likelihood of a doublet/multiplet by creating a subset of artificial “doublets” by merging two individual cells together [58]. Theoretically, this method will detect doublets composed of two different cell types, however, since the vast majority (∼70%) of cells captured were oligodendrocytes, this means that the vast majority of doublets will be largely undetectable. However, with a cell capture of 8,000 cells per sample, 10x estimates a 6.4% multiplet rate. Clusters lacking uniquely expressed genes (and thus a unique cell identity) were excluded from downstream analyses as they were most likely multiplets of two different cell types. All scripts used in this study are available on our GitHub page (https://github.com/UW-BRaIN-lab).

### Dorsolateral Pre-Frontal Cortex Gray Matter Dataset

The DLPFC GM single nucleus RNA sequencing dataset was obtained from the Allen Institute’s Seattle Alzheimer’s Disease Brain Cell Atlas (SEA-A). All 80 DLPFC cases were downloaded as .h5ad files converted to Seurat objects using sceasy (https://github.com/cellgeni/sceasy) and the reticulate (https://rstudio.github.io/reticulate) R packages. Once converted to Seurat objects, all files were merged and metadata aligned to the existing WM dataset. The SEA-AD dataset included 18 donors (9 high ADNC, 9 no ADNC) from the WM dataset developed, as well as 68 other donors (this cohort’s summary statistics are included in Table 1). Generally, the SEA-AD dataset was processed in the same manner as the WM cohort, using 10x v3.1 GEM kits, sequence alignment and mapping through CellRanger (mapped to Human genome assembly GRCh38), and similar thresholds for low-quality cell quality control (i.e. minimum of 500 unique genes per cell). However, there are a few differences between our methods. First, SEA-AD sought to sequence to a depth of 120,000 reads per cell, whereas our WM dataset was sequenced at 40,000 reads per cell. Both of these values are above the 10x-recommended 20,000 reads per cell minimum. Second, SEA-AD employed FACS sorting to load 10,000 cells per well (compared to 8,000 for the WM) at a ratio of 70% NeuN+/30% NeuN- cells through 10x, enriching for neurons at the expense of the non-neural cell types. This means percent representation of each glial cell type cannot be directly compared between WM and GM. Leveraging the Allen Institute’s prowess in characterizing and classifying cell types, all “supertypes” of each subclass (Oligodendrocyte, OPC, etc.) of the DLPFC dataset were subset into individual Seurat objects to be merged with the WM.

### Separation of WM clusters and merger with DLPFC GM clusters by cell type

Individual sample Seurat objects were merged and integrated using canonical correlation analysis (CCA), which is documented here[82]. This process resulted in the UMAP shown in Figure 1. Each cluster’s general cell type was identified using unique identification genes, primarily based on previous cell mapping done by the Allen Institute (https://portal.brain-map.org/explore/seattle-alzheimers-disease). At this point, clusters were isolated into individual Seurat objects depicting each cell type which were used for downstream analysis. Clusters that contained 5 or less cells were excluded from downstream analysis. The WM and DLPFC GM individual cell type Seurat objects were merged and reanalyzed using the standard Seurat pipeline (normalize, FindVariableFeatures (nFeatures=5,000 or 10,000), scale, PCA, UMAP (dimensions 1:15-1:20 depending on ElbowPlot, resolution of clusters optimized to return between 5-10 clusters for OLs).

### NanoString CosMx Spatial Transcriptomics 6,000-plex Human Discovery Panel Experimental Design and Procedure

Six slides (three runs of two slides) were processed and ran through the NanoString/Bruker CosMx single cell spatial platform, which uses a combination of in-situ hybridization probes and UV cleavable fluorescent dyes to accurately label and identify transcripts at single-cell resolution level. This technology includes negative probes to address non-specific binding and NanoString-developed initial data analysis software (AtoMx).

Of the 18 donors that were analyzed both in our WM cohort and in SEA-AD, we selected twelve to analyze using CosMx. These donors were separated by sex, where each slide contained one male/female with high AD pathology (ADNC=3) and one male/female with not/low AD pathology (ADNC=0/1). In addition to having each slide be male or female, we also included a fresh frozen white matter and DLPFC gray matter sample from each donor. Thus, each slide had a total of four samples, 2 WM and 2GM. Tissue samples were obtained following the same guidelines as described above for the 10x process, with the WM samples being the immediate adjacent WM to the 10x core and the DLPFC GM cores being obtained using images provided by the Allen Institute to ensure the region dissected was as close to the SEA-AD sample as possible. Following dissection, samples were transferred to a cryostat and aggregate blocks were assembled in Optimal-Cutting Temperature compound (OCT) rapidly to avoid freeze-thawing. Blocks were sectioned at 5um and slides stored at -80C until CosMx procedure.

The CosMx procedure using fresh frozen tissue was followed (MAN-10184-03, September 2024 publish date), using the “CosMx Human Universal Cell Segmentation Kit (RNA) 1.0), which includes anti-human ribosomal RNA, Histone H3, and GFAP markers in addition to DAPI. The “CosMx Human 6K Discovery Panel, 6K-Plex, RNA” probe set was used (Item #: 121500041). Following completion of slide preparation, slides were loaded onto the machine and run initiated following “CosMx SMI Instrument User Manual (software v1.3)” (MAN-10161-05, published February 2024). A maximum of 400 fields of view were selected for each two-slide run, 200 per slide, to ensure tissue integrity was maintained throughout entire run. WM FOVs were selected at random throughout each sample, whereas GM FOVs were selected to span the entire cortical ribbon to ensure any regional variability contained within the 10x data (which included the entire cortical ribbon) would be also present in the CosMx data. Following data collection, raw data is autonomously processed and demultiplexed on NanoString’s AtoMx cloud-based analysis software. Cell segmentation was sufficient using standard NanoString recommendations, specifically pre-bleaching profile Configuration C (60 seconds), cell segmentation Configuration B, and exposure times and thresholding as default for each morphology marker. This provided reasonable cell segmentation masks which were then carried forward to QC and data analysis.

### CosMx Quality Control and Data Analysis

Following data acquisition, a study was created in AtoMx to provide an initial analysis of this data. The AtoMx-provided pipeline “Foundational modules (RNA)” was executed on the dataset with default NanoString-provided settings. This pipeline is largely based on the Seurat pipeline described above, with the addition of a unique NanoString-developed cell type identification algorithm known as InSituCellTypes (https://github.com/Nanostring-Biostats/InSituType). This algorithm uses a semi-supervised clustering method where an unknown dataset is mapped onto an existing and annotated dataset to identify cell types based on established identifiers. Importantly, any cells that do not share similarity with any of the provided profiles will be clustered into separate groups for further study. For this study, we mapped our CosMx WM and GM data together using the semi-supervised method, mapping to the NanoString-provided human brain cell profiles (https://github.com/Nanostring-Biostats/CosMx-Cell-Profiles/tree/main/Human/Brain). This brain profile dataset was developed using human formalin-fixed, paraffin-embedded frontal gray matter processed using the same “CosMx Human 6K Discovery Panel, 6K-Plex, RNA” probe set as our study, and thus was the most applicable reference to map our data to.

After completion of this pipeline, the dataset was exported as Seurat objects to be further analyzed in R. The exported objects included all dimensional reduction and cluster information derived from the AtoMx pipeline. The bimodal distribution of our clusters on UMAP (see Fig.2A) suggested there was some technical aspect that was causing the dimensional reduction to segregate each cluster into two partitions. We identified that the AtoMx QC process flagged cells but did not remove them, so we proceeded to implement our own QC. First, we removed any FOV with a mean reads/cell of 300 or less, indicative of low-quality FOVs. Next, we removed any cells with less than 200 reads/cell. Finally, we removed the top 6% of cells by area (area > 30,000um^2^) under the assumption that these “cells” were likely multiple cells segmented as one or another segmentation artifact. Following this QC process, all clusters identified in InSituCellTypes() were still represented, but as expected, the majority of one branch of the bimodal UMAP was removed.

There was a clear batch effect between individual runs, so we corrected this using the same Seurat canonical correlation analysis (CCA) used for our 10x data. We ran the entire pipeline we developed for our 10x data on the CosMx data, with the only change being adjusting the resolution of the clustering after integration to merge some of the cell type subclusters. This CCA integration seemed to align all individual slides well, and provided concrete and unique cell type clusters that were used for downstream analyses.

### Differential Expression, Pathway Analysis, Data Visualization

These analyses were conducted on both the 10x-derived WM and GM cell types as well as the CosMx cell types. Differential expression analyses were conducted using FindMarkers() in Seurat, with the logfc threshold set to zero, and percent of cells expressing each transcript set to 1% (unless otherwise noted), thus analyzing every cell and nearly all genes in the dataset. Pathway analysis/gene enrichment was conducted via the R package “gage” (v2.54.0), using the Human EnsDb.Hsapiens.v86 (v.2.99.0) database to map gene identifications. Both Gene Ontology (GO) and Kyoto Encyclopedia of Genes and Genomes (KEGG) were assessed in pathway analysis. Log2fc is the default method for gage() and we decided to use that for our analysis. Second, the q-values were reported, as these were adjusted p-values that account for the false discovery rate using the Benjamini-Hochberg method. Figures were generated using RStudio’s ggplot2 (v3.5.1) with error bars, color coordination, and formatting done in Adobe Illustrator 2024. All datasets are available for download at on Zenodo at DOI: 10.5281/zenodo.22133455.

### Statistics

For differential expression testing, p-values were calculated autonomously as part of the Seurat FindMarkers() and adjusted using the Benjamini-Hochberg correction method. In figures, these p-values are depicted as inverted base-10 logarithms with dotted lines denoting an adjusted p-value of 0.05 to indicate significance threshold. For pathway analyses, the log2fc values calculated during differential expression were used as input. Similar to the differential expression analysis, mean log2fc and adjusted p-values using the Benjamini-Hochberg method were reported. For assessing significance of cluster representation between the various groups (Young, No AD, and AD), the representation of each cell type (as a percentage of total cells captured in that sample) was calculated. These percentages were then used to generate the resulting cluster by cluster distributions. Significance was assessed via one-way ANOVA with Tukey’s HSD test.

### Pseudotime inference in oligodendroglia

Pseudotime was computed across the oligodendroglial lineage using diffusion pseudotime (DPT) implemented in Scanpy. Briefly, oligodendroglial states were defined from the Supertype_fixed annotation and renamed for clarity (NFOL/differentiating, lipid remodeling, mature myelinating, and stress/ISR-reactive). To ensure stable trajectory estimation, DPT was restricted to states containing at least 5,000 cells; cells outside these states were not assigned pseudotime. To reduce computational burden, a random subsample of 90,000 eligible cells was used to compute DPT. A k-nearest-neighbor graph was constructed on the integrated low-dimensional embedding (PCA coordinates stored in X_pca, derived from the Seurat CCA-integrated dataset) using 30 neighbors. DPT was rooted in NFOL/differentiating oligodendrocytes (BCAS1/ENPP6+) by selecting a representative NFOL cell from the subsample as the root (iroot). DPT pseudotime values computed on the subsample were then projected to all eligible cells by kNN regression in the same embedding (k = 15), using an inverse-distance weighted average of the subsample pseudotime values.

### Cell state transitions

We quantified directed state transitions using a kNN graph constructed in the same integrated low-dimensional embedding used for pseudotime (PCA coordinates from the Seurat CCA-integrated dataset). For each cell, we evaluated its k nearest neighbors and counted “forward” edges to neighbors with an increase in DPT pseudotime of at least Δt ≥ 0.03. For each source state A, forward neighbor counts were aggregated into target states B and normalized by the total number of forward edges exiting A to yield a directed transition weight *P_dir_*(*A* ➔ *B*), interpreted as the fraction of pseudotime-forward edges from A that terminate in B. Networks display edges with *P_dir_*: 2 0.04; node size reflects the number of cells per state and edge width reflects *P_dir_*.

